# Postsynaptic BMP signaling regulates myonuclear properties in *Drosophila* larval muscles

**DOI:** 10.1101/2024.04.10.588944

**Authors:** Victoria E. von Saucken, Stefanie E. Windner, Mary K. Baylies

**Affiliations:** Developmental Biology Program, Sloan Kettering Institute, Memorial Sloan Kettering Cancer Center, New York, NY 10065 USA; Weill Cornell-Rockefeller-Sloan Kettering Tri-Institutional MD-PhD Program, New York, NY 10065 USA; Biochemistry, Cell & Developmental Biology, and Molecular Biology (BCMB) Program, Weill Cornell Graduate School of Medical Sciences, New York, NY 10065 USA

**Keywords:** nuclear size scaling, pMad, endoreplication, neuromuscular junction, myonuclear domain, synaptic nuclei

## Abstract

The syncytial mammalian muscle fiber contains a heterogeneous population of (myo)nuclei. At the neuromuscular junction (NMJ), myonuclei have specialized positioning and gene expression. However, it remains unclear how myonuclei are recruited and what regulates myonuclear output at the NMJ. Here, we identify specific properties of myonuclei located near the *Drosophila* larval NMJ. These synaptic myonuclei have increased size in relation to their surrounding cytoplasmic domain (scaling), increased DNA content (ploidy), and increased levels of transcription factor pMad, a readout for BMP signaling activity. Our genetic manipulations show local BMP signaling affects muscle size, nuclear size, ploidy, and NMJ size and function. In support, RNA sequencing analysis reveals that pMad regulates genes involved in muscle growth, ploidy (i.e., *E2f1*), and neurotransmission. Our data suggest that muscle BMP signaling instructs synaptic myonuclear output that then positively shapes the NMJ synapse. This study deepens our understanding of how myonuclear heterogeneity supports local signaling demands to fine tune cellular function and NMJ activity.

**Summary:** The neuromuscular junction (NMJ) is a well characterized synapse, yet the postsynaptic contributions that allow for synapse function are not well understood. This study by von Saucken *et al*. uses the *Drosophila* larval NMJ to define synaptic muscle (myo)nuclei and their properties and determine how BMP signaling regulates these myonuclear properties.

## Introduction

Skeletal muscle cells (myofibers) contain hundreds of post-mitotic (myo)nuclei that exhibit heterogeneity despite sharing one cytoplasm. Single nuclear sequencing in mammalian muscle revealed that myonuclei have distinct transcriptional profiles based on their intracellular position (Dos Santos et al., 2020; Kim et al., 2020; Petrany et al., 2020). For example, myonuclei adjacent to the neuromuscular junction (NMJ) express genes specifically required for the myofiber’s postsynaptic structures. These intracellular differences are consistent with the myonuclear domain hypothesis that considers the limited synthetic capacity of individual myonuclei and the physical constraints on cellular transport and diffusion (Hall and Ralston, 1989; Pavlath et al., 1989). It proposes that each nucleus within the syncytial myofiber supplies gene products only to its immediately surrounding cytoplasm (myonuclear domain). Thus, myonuclei are spaced throughout the myofiber to allow for efficient distribution of gene products for muscle homeostasis and function. In contrast, myonuclei adjacent to the NMJ are clustered, which locally increases nuclear size scaling (relationship between myonuclear size and surrounding cytoplasmic domain size) and DNA content (Sanes and Lichtman, 2001), resulting in local enrichment of NMJ components at the postsynaptic membrane (Duclert and Changeux, 1995; Schaeffer et al., 2001). An open question is how synaptic myonuclei are specified and their activity programmed to provide gene products for effective synaptic transmission.

The NMJ is a specialized synapse composed of the myofiber’s postsynaptic structures in opposition to the motor neuron’s presynaptic axonal endings (boutons). The *Drosophila melanogaster* larva provides an easily accessible and a highly conserved NMJ, ideal for studying synaptic development and function through genetic manipulations and visualization (Broadie and Bate, 1995; Deng et al., 2017). While presynaptic bouton formation and maturation have been thoroughly examined (reviewed in Chou et al., 2020), the postsynapse is far less studied. The *Drosophila* postsynapse consists of complex membrane folds (subsynaptic reticulum, SSR) that increase muscle surface area. Membrane receptors within the SSR, including glutamatergic neurotransmitter receptors, participate in synaptic signaling which leads to muscle contraction (DiAntonio et al., 1999; Nguyen and Stewart, 2016; Zou and Pan, 2022). The SSR develops following new bouton formation (Vasin et al., 2019), and critical regulators of SSR development include scaffolding proteins (Lahey et al., 1994), adhesion proteins (Banovic et al., 2010; Xing et al., 2018), and actin modulators (Blunk et al., 2014; Christophers et al., 2023). Like at the vertebrate postsynapse, *Drosophila* myonuclei can support SSR development. For example, work on synapse-to-myonucleus signaling revealed that the Frizzled nuclear import pathway promotes SSR maturation, but the mechanism is not fully understood (Restrepo et al., 2022; Speese et al., 2012). However, it is unknown whether all *Drosophila* myonuclei are equally affected by NMJ-derived signals and/or establish transcriptional heterogeneity. Previous work from our lab and others has identified differences in myonuclear position, size scaling, and DNA ploidy in the center of the muscle cell near the larval NMJ (Perillo and Folker, 2018; Windner et al., 2019). Nevertheless, *Drosophila* synaptic myonuclei remain uncharacterized, providing an opportunity to advance our understanding of postsynaptic adaptions and investigate whether local myonuclear changes are a conserved muscle feature.

Bone morphogenetic protein (BMP) ligands are critical signaling proteins at both the mammalian and *Drosophila* NMJ (Bayat et al., 2011; Fish et al., 2023; Yilmaz et al., 2016). In *Drosophila*, a retrograde muscle-to-motor neuron BMP pathway that promotes NMJ growth has been extensively studied (Aberle et al., 2002; Marqués et al., 2002; McCabe et al., 2003). The BMP ligand, Glass bottom boat (Gbb), is secreted by the postsynaptic muscle and activates the presynaptic neuronal BMP receptors. This leads to phosphorylation and nuclear import of the transcriptional effector Mad (pMad), which regulates expression of key presynaptic NMJ genes (Ball et al., 2010; Kim and Marqués, 2010; Smith et al., 2012). Studies in BMP signaling mutants revealed that loss of BMP signaling results in reduced bouton number and impaired NMJ synaptic transmission, specifically reduced release of neurotransmitters and a diminished postsynaptic excitatory response (Aberle et al., 2002; Marqués et al., 2002; McCabe et al., 2003). These effects are attributed to disrupting the retrograde pathway because transgene expression of BMP effectors specifically in the motor neuron, but not in the muscle cell, rescued these mutant NMJ defects (Aberle et al., 2002; McCabe et al., 2004).

However, some data suggest that postsynaptic BMP signaling contributes to NMJ development and muscle growth. BMP signaling mutants have smaller muscles, which are rescued to normal size by muscle-specific expression and only partially rescued by neuronal expression (McCabe et al., 2004). The muscle expresses BMP receptors and Mad, which localize to the postsynaptic membrane, and the active form of Mad (pMad) is reduced at the NMJ following muscle-specific Mad knockdown (KD) (Dudu et al., 2006; Fuentes-Medel et al., 2012). Like the retrograde pathway, manipulations of postsynaptic BMP signaling affect NMJ structure and neurotransmission (Fuentes-Medel et al., 2012; Rawson et al., 2003; Sulkowski et al., 2016). However, it remains to be investigated whether postsynaptic BMP signaling affects the morphology, position, and transcriptional output of myonuclei.

In this study, we define synaptic myonuclei in *Drosophila* larval muscles and find that they are distinct in size and DNA content. Further, we find that synaptic myonuclei are enriched in pMad, the active BMP transcription factor. Unlike previous work, we investigate the effects of postsynaptic BMP signaling on synaptic myonuclei by muscle-specific KD of the core BMP signaling components: the type II receptor Punt, the type I receptor Thickveins, and Mad. We show that postsynaptic BMP signaling regulates muscle size, nuclear size, and DNA ploidy. RNA sequencing reveals KD of Mad reduces mRNA levels of the critical G1/S regulator *E2f1*, suggesting that postsynaptic BMP signaling promotes nuclear DNA content by regulating the endoreplication cell cycle program. We also find changes in the expression of NMJ genes (i.e., Glutamate receptor subunits) and in NMJ structure and function in Mad KD muscles. We propose that local BMP signaling at the postsynapse exposes synaptic myonuclei to higher levels of pMad, which in turn, result in localized increased endoreplication and expression of NMJ-related genes that influence synaptic structure and function. This study provides novel insights to the mechanisms promoting myonuclear heterogeneity and postsynaptic development and has importance for our understanding of processes underlying NMJ functionality in the contexts of development and disease.

## Results

### Defining synaptic myonuclei in *Drosophila* larval muscles

To characterize synaptic myonuclei at the *Drosophila* NMJ, we used the well-studied ventral longitudinal muscles VL3 and VL4 (muscles 6 and 7) from abdominal segments 2 and 3 (Fig. 1A). VL3 and VL4 muscles differ in size, nuclear number, and nuclear arrangement in two rows versus one row (Fig. S1A-B; Manhart et al., 2018; Windner et al., 2019). While the innervating motor neuron is shared between VL3 and VL4, each muscle develops its own NMJ (Kohsaka et al., 2012). We previously identified a nuclear size scaling relationship between myonuclei and their surrounding cytoplasmic domain. Both nuclear size scaling and DNA content were found to be intracellularly patterned: myonuclei with the largest size and highest DNA ploidy occur in the middle of the muscle cell where the NMJ is located (Windner et al., 2019). Here we asked whether myonuclei closer to the NMJ are significantly different from those positioned more distant to the NMJ.

**Figure 1.**
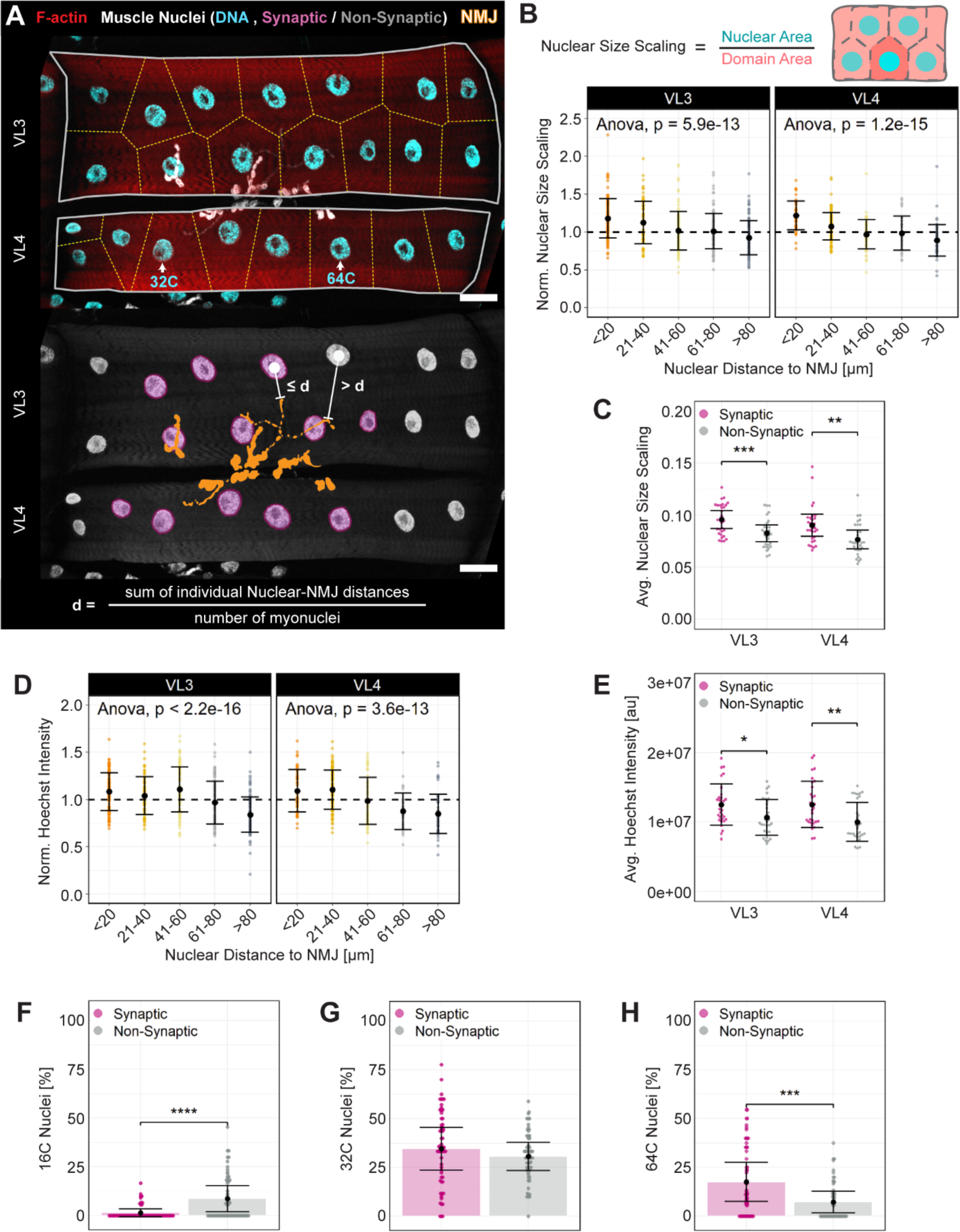
Synaptic myonuclei in *Drosophila* larval muscles are characterized by position, size, and DNA content. **(A)** Confocal images (sum slice projection) of ventral longitudinal muscles VL3 and VL4 (or muscles 6 and 7) from wandering *Dmef2>mCherry-RNAi* control larvae. **Top**, fluorescent labeling of muscle cells (phalloidin, red), nuclear DNA (Hoechst, cyan), and NMJ (anti-Dlg, white). Myonuclear domains (yellow dashed lines) and example myonuclei with ploidy of 64C and 32C are indicated. **Bottom**, same cells highlighting the NMJ (orange), NMJ-proximal synaptic myonuclei (magenta), and distal non-synaptic myonuclei (dark gray). Synaptic myonuclei are defined per cell as nuclei positioned less than or equal to the average nuclear distance to the NMJ (d). Scale bars = 30 µm. **(B)** Nuclear size scaling, equal to nuclear size divided by cytoplasmic domain size, normalized by the cell’s average and plotted against distance from the NMJ (20 µm bins). **(C)** Average nuclear size scaling for synaptic and non-synaptic myonuclei. **(D)** Hoechst fluorescence intensity, normalized by the cell’s average and plotted against distance from the NMJ (20 µm bins). **(E)** Average Hoechst fluorescence intensity for synaptic and non-synaptic myonuclei. **(F-H)** Ploidy distribution based on Hoechst fluorescence intensities (refer to Fig. S1H) in synaptic and non-synaptic myonuclei from VL3 and VL4 muscles for **(F)** 16C, **(G)** 32C, and **(H)** 64C nuclei. Data shown as mean ± SEM, except for in B and D where mean ± SD is plotted (*n =* 19 larvae, 30 VL3 muscles and 30 VL4 muscles). Statistical comparisons were made for all groups with significant differences noted (* p < 0.05, ** p < 0.01, *** p < 0.001, **** p < 0.0001).

To identify the NMJ, we immunolabeled for the scaffolding protein Discs large (Dlg) that closely associates with the SSR and is a widely used postsynaptic marker (Chen and Featherstone, 2005; Zhang et al., 2017). We measured the distance between the nuclear centroid and the nearest Dlg-positive NMJ element (Fig. 1A, S1C). Since synaptic myonuclei cannot be identified visually, we used the average distance between myonuclei and the NMJ (d) to separate the myonuclei into two populations, synaptic nuclei (≤ d) and non-synaptic nuclei (> d). The average nuclear-NMJ distance was 56.25 ± 9.98 µm for the larger VL3 muscle and 49.23 ± 8.87 µm for the smaller VL4 muscle (mean ± SD), with a mean of 8 synaptic nuclei (∼48% of nuclei) in VL3 and 6 synaptic nuclei (∼58% of nuclei) in VL4 (Fig. S1D-E). Given these differences, we initially examined myonuclear attributes separately for VL3 and VL4 muscles.

Increased myonuclear size scaling at the mammalian NMJ indicates an increased nuclear potential to contribute gene products to the NMJ. To assess nuclear size scaling, we measured the size of nuclei and their cytoplasmic domains as previously described (Windner et al., 2019). We found that nuclear size scaling is significantly increased in myonuclei closer to the NMJ, dropping below average for myonuclei at ∼41-60 µm (diameter of approximately 3 nuclei) distance (Fig. 1B). This size scaling increase resulted from an increase in myonuclear sizes, while cytoplasmic domain sizes remained unchanged (Fig. S1F-G). When calculating average nuclear size scaling for synaptic and non-synaptic myonuclei, we found similar values in both VL muscles (Fig. 1C). Therefore, in both VL3 and VL4, synaptic myonuclei have significantly increased nuclear size scaling compared to non-synaptic myonuclei.

Nuclear ploidy correlates with cell size and polyploidy functions to support growth and the metabolic demands of large cells (Edgar and Orr-Weaver, 2001). In addition to each larval muscle containing multiple nuclei, each myonucleus increases its DNA content by endoreplication, resulting in copy numbers of 16C, 32C, and 64C (Windner et al., 2019). To assess myonuclear DNA content, we used the fluorescence intensity of the DNA stain Hoechst, as done previously (Windner et al., 2019). We found, similar to nuclear size scaling, that myonuclei with increased Hoechst intensity were positioned less than or equal to 60 µm from the NMJ, supporting the use of average nuclear-NMJ distance for identifying synaptic myonuclei (Fig. 1D). We calculated average Hoechst intensity for synaptic and non-synaptic myonuclei, revealing that synaptic myonuclei have significantly increased DNA content in both VL muscles (Fig. 1E). To further evaluate nuclear DNA content, we assigned ploidy numbers to Hoechst intensity measurements (Fig. S1H). This analysis revealed that synaptic myonuclei have higher ploidy numbers (more 64C nuclei and fewer 16C nuclei) (Fig. 1F-H), indicating that they undergo additional rounds of endoreplication compared to non-synaptic myonuclei.

At the mammalian NMJ, increased DNA content and nuclear size scaling are achieved by clustering myonuclei at the postsynapse (Bruusgaard et al., 2003; Hansson et al., 2020). We observed no change in cytoplasmic domain sizes for myonuclei near the *Drosophila* NMJ (Fig. S1G). To directly test whether clustering occurs, we measured internuclear distance and observed similar distances for synaptic and non-synaptic myonuclei in VL3 and VL4 muscles (Fig. S1I). Thus, at the *Drosophila* NMJ, a local increase in myonuclear DNA content and nuclear size scaling is established by providing more DNA per synaptic nucleus and increasing nuclear size, respectively, in the absence of nuclear clustering. These synaptic nuclear properties are observed in both VL muscles, indicating that it is a general feature of these larval muscles. Based on these data, we define populations of synaptic and non-synaptic myonuclei in *Drosophila* muscles by their distance to the NMJ, nuclear size scaling, and DNA content.

### Synaptic myonuclei have increased BMP signaling

A key signaling pathway at the NMJ is the bone morphogenetic protein (BMP) pathway. While postsynaptic pathway activation and nuclear import have been described (Dudu et al., 2006), how this affects the postsynaptic muscle, particularly the myonuclei, is unknown. To confirm that the core BMP pathway components (Fig. 2A) are expressed in larval muscles, we analyzed publicly available RNA sequencing data of larval muscle-enriched preps (Christophers et al., 2023). We found moderate expression of genes encoding the BMP type II receptor *put* (Punt), type I receptors *tkv* (Thickveins) and *sax* (Saxophone), BMP transcriptional effector *Mad* and its co-factor *Med*, and BMP ligand *gbb* (Glass bottom boat) (Fig. 2B). Interestingly, *tkv* was more highly expressed than the other type I receptor *sax*. This suggests that, similar to BMP signaling in *Drosophila* wing development (Haerry et al., 1998), a preferential use of type I receptor Tkv over Sax occurs in larval muscle.

**Figure 2.**
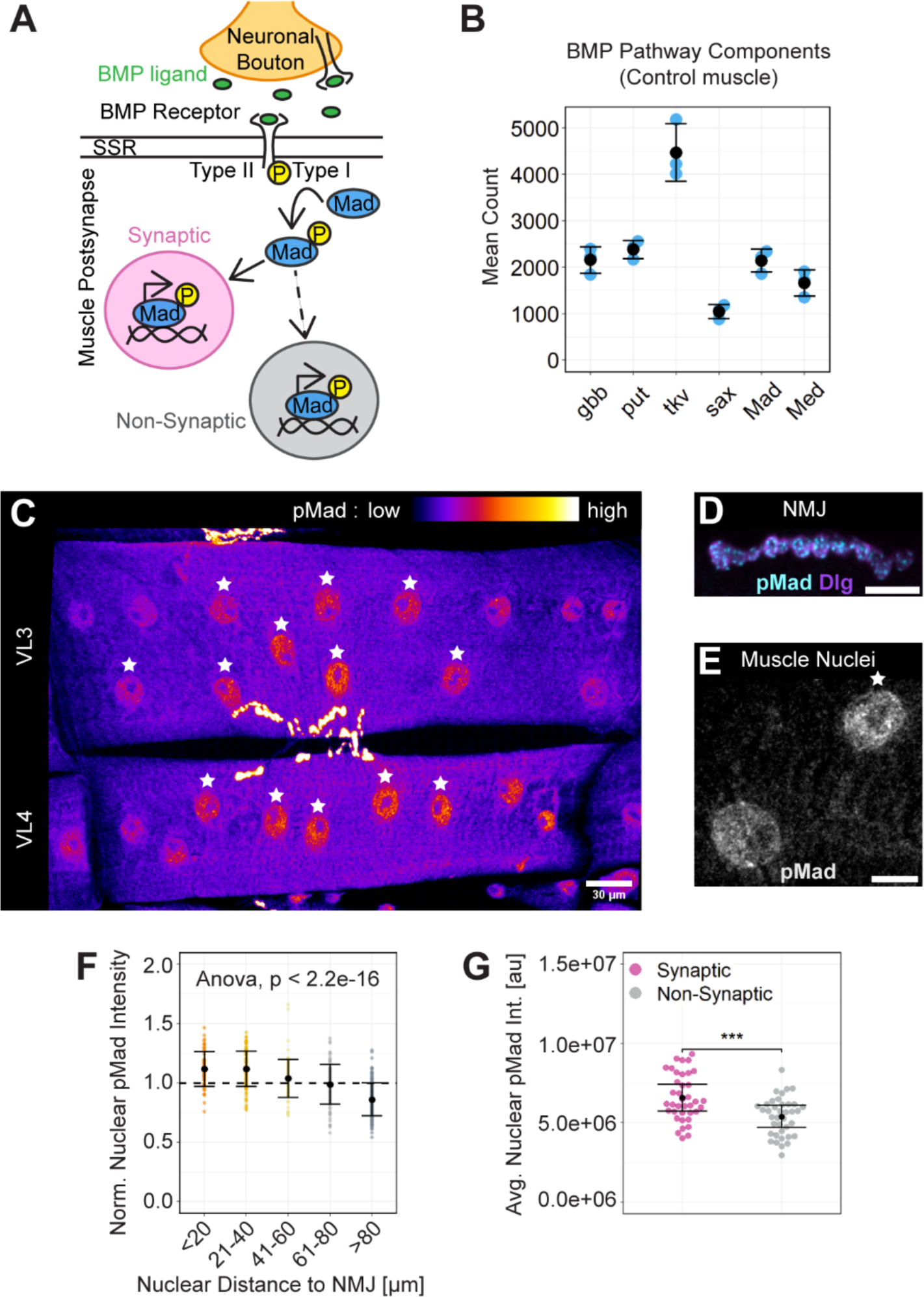
BMP signaling effector pMad is increased within synaptic myonuclei. **(A)** Diagram of the postsynaptic BMP signaling pathway in *Drosophila* larval muscle. BMP ligand (Gbb, green) binds to receptor hetero-tetramers composed of two Type I (Tkv or Sax) and two Type II (Punt) receptors in the postsynaptic muscle membrane (SSR) opposed to the neuronal boutons (orange). Activation of the receptor complex leads to phosphorylation of Mad (pMad, blue), its nuclear translocation, and subsequent regulation of target gene transcription. We hypothesize that synaptic myonuclei (magenta) are exposed to higher pMad levels than non-synaptic myonuclei (gray). **(B)** Normalized mRNA counts for *Drosophila* BMP pathway genes in third instar control *Mhc>mCherry-RNAi* larvae from a published bulk RNA sequencing dataset (Christophers et al., 2023). **(C)** Confocal image (sum slice projection) of VL3-VL4 muscle pair from a *Dmef2>mCherry-RNAi* control larva immunolabeled for pMad. Colors indicate pixel intensities, with highest levels at the NMJ (white-yellow) and within synaptic myonuclei (white stars). **(D)** Confocal image of a VL3/4 NMJ branch (sum slice projection) showing pMad (cyan) and SSR (Dlg, magenta). **(E)** Confocal image of nuclear pMad (sum slice projection) in a synaptic nucleus (white star) next to a non-synaptic nucleus. Scale bars = 30 µm (C), 10 µm (D, E). **(F)** Nuclear pMad intensity, normalized by the cell’s average and plotted against distance from the NMJ (20 µm bins) (*n =* 11 larvae, 40 muscle pairs of VL3 and VL4). **(G)** Average nuclear pMad intensity in synaptic and non-synaptic myonuclei from control muscles (*n =* 11 larvae, 40 muscle pairs of VL3 and VL4). Data shown as mean ± SEM in plot G and as mean ± SD in plots B and F (*** p < 0.001).

To confirm BMP signaling activity in *Drosophila* larval muscles, we performed immunolabeling for pMad. We observed strong, punctate pMad signal directly at the NMJ as well as diffuse and punctate pMad signal in myonuclei (Fig. 2C-E). These patterns are consistent with previous studies (Dudu et al., 2006; O’Connor-Giles et al., 2008; Smith et al., 2012; Sulkowski et al., 2014). When we analyzed the distribution of nuclear pMad signal, we found that the pMad signal is significantly increased in myonuclei closer to the NMJ, dropping below average in myonuclei greater than 60 µm from the NMJ (Fig. 2F). We calculated average nuclear pMad intensity, revealing that synaptic myonuclei have greater pMad signal than non-synaptic myonuclei (Fig. 2G), in both VL muscles (Fig. S2A). This increase remained when nuclear pMad intensity was normalized to nuclear area, indicating that synaptic myonuclei demonstrate higher pMad levels independent of size (Fig. S2B). Thus, BMP pathway activation originating from the NMJ establishes a gradient across the myonuclei and high levels of pMad are characteristic of the synaptic nuclear population.

### Postsynaptic BMP signaling promotes muscle size and myonuclear size

To explore the effects of postsynaptic BMP signaling on synaptic myonuclei, we performed muscle-specific, RNAi-mediated knockdown (KD) of two BMP receptors, Punt (type II) and Tkv (type I), and the downstream transcriptional effector Mad (Fig. 3A). We determined BMP target KD efficiencies by qPCR on larval muscle-enriched preps, revealing a 20-30% KD of *put*, *tkv*, and *Mad* levels (Fig. S2C). Given the similar nuclear patterns in VL3 and VL4 control muscles, we combined data from the VL muscles for the analysis of KD muscles. Muscle KD of Punt, Tkv, or Mad showed a reduction in the size of both muscle cells and myonuclei (Fig. 3B; S2D), indicating a role for postsynaptic BMP signaling in promoting muscle growth. Reduction of both cell and nuclear sizes resulted in similar nuclear size scaling for synaptic and non-synaptic myonuclei in Tkv KD, Punt KD, and control muscles. However, in Mad KD muscles, only non-synaptic myonuclei had control-like size scaling, while that of synaptic myonuclei was reduced (Fig. 3C). These data suggest that Mad KD results in a stronger phenotype than receptor level KD and more strongly affects synaptic myonuclei. Despite overall reduced sizes, muscles of all genotypes, maintain increased nuclear size scaling for synaptic relative to non-synaptic myonuclei.

**Figure 3.**
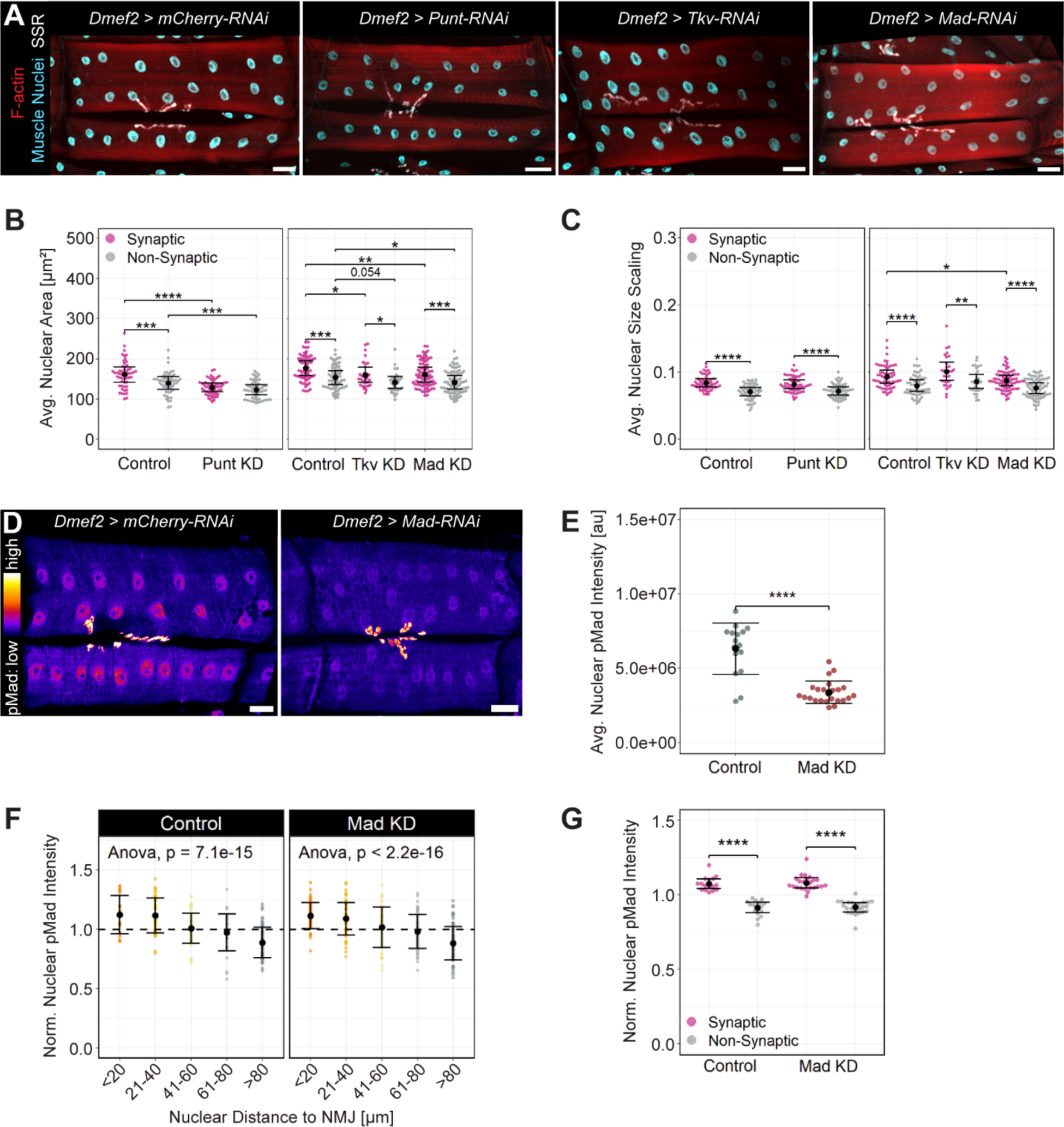
Muscle-specific KD of BMP pathway components reduces nuclear size and pMad levels but maintains a nuclear pMad gradient. **(A)** Confocal images of VL muscles (sum slice projection) labeled with phalloidin (F-actin, red), Hoechst (DNA, cyan), and anti-Dlg (SSR, white) in control and Punt, Thickveins (Tkv), and Mad KD (KD) muscles. Scale bars = 30 µm. **(B)** Average size of synaptic and non-synaptic myonuclei in control and Punt-deficient VL muscles (*n =* 17 larvae, 56 muscles in both groups), and control, Tkv, and Mad KD muscles (control: *n =* 19 larvae, 60 muscles; Tkv KD *n =* 12 larvae, 30 muscles; Mad KD *n =* 20 larvae, 76 muscles). **(C)** Average nuclear size scaling for synaptic and non-synaptic myonuclei. Genotypes and sample numbers as in (B). **(D)** Confocal images of control and Mad KD muscles (sum slice projection pseudo-colored to indicate pixel intensity) labeled for pMad. Scale bars = 30 µm. **(E)** Average nuclear pMad intensity in control and Mad KD muscles (control: *n =* 6 larvae, 16 muscles; Mad KD: *n =* 9 larvae, 24 muscles). **(F, G)** Nuclear pMad intensity, normalized by the cell’s average, **(F)** plotted against distance from the NMJ (20 µm bins), and **(G)** shown for synaptic and non-synaptic myonuclei in control and Mad KD muscles. Data shown in all plots as mean ± SEM, except in F where mean ± SD is plotted. Statistical comparisons were made for all groups with significant differences noted (* p < 0.05, ** p < 0.01, *** p < 0.001, **** p < 0.0001).

Muscle KD of Punt, Tkv, or Mad showed average nuclear distances to the NMJ similar to controls, despite smaller muscles (Fig. S2E). We attribute this to a decrease in relative NMJ length, not a difference in nuclear distribution (Fig. S2F). In accordance, the percent of synaptic myonuclei per muscle was unchanged in most groups. However, VL3 muscles deficient in Tkv and Mad had a ∼9-14% increase in synaptic myonuclei compared to control VL3 muscles (Fig. S2G). These observations indicate that *Drosophila* muscles can adjust nuclear positioning in relation to the NMJ to compensate for limited growth.

Next, we assessed nuclear pMad levels in Mad KD muscles, which revealed that average protein levels are reduced by ∼50% (Fig. 3D-E). When we examined the intracellular distribution of nuclear pMad, we found that Mad KD muscles maintained a nuclear pMad gradient as observed in control muscles, as well as significant differences between synaptic and non-synaptic nuclei (Fig. 3F-G). Together, these results indicate that perturbations at the BMP receptor and transcriptional effector levels affect muscle growth, and additionally in Mad KD muscles, reduced nuclear size scaling of synaptic myonuclei. Nevertheless, the patterns of nuclear size scaling (synaptic vs. non-synaptic) as well as the pMad gradient are maintained as in controls.

### Postsynaptic BMP signaling promotes myonuclear DNA content via regulation of E2f1

We next determined whether postsynaptic BMP signaling affects myonuclear DNA content (Fig. 4A). We observed a strong positive correlation between total Hoechst intensity and total nuclear pMad intensity in control muscles and in Mad KD muscles, supporting a relationship between postsynaptic BMP signaling and myonuclear DNA content (Fig. 4B). We found that average nuclear Hoechst intensity was reduced in Mad KD and Punt KD muscles, but similar to controls in Tkv KD muscles (Fig. 4C). Further, average nuclear Hoechst intensity is increased in synaptic compared to non-synaptic nuclei in Tkv KD and Mad KD muscles; however, no significant differences were observed in Punt KD muscles. To achieve a more detailed analysis of DNA content, we used Hoechst intensity to designate nuclear ploidy (Fig. S3A). In control muscles, synaptic nuclei exhibit increased ploidy (more 64C and less 16C) compared to non-synaptic nuclei. Consistent with average Hoechst intensity, overall 64C ploidy was reduced in muscles with KD of Punt and Mad, but not Tkv (Fig. 4D-F). Muscles across all groups maintained significantly higher ploidy (greater 32C and/or 64C) in the synaptic myonuclei, revealing that a local DNA increase is maintained at the postsynapse. A reduction of 64C nuclei in Punt KD and Mad KD muscles correlated with a significant increase of 32C ploidy in the synaptic nuclei compared to the non-synaptic nuclei (Fig. 4D, 4F). These data reveal that perturbations to BMP signaling affect nuclear DNA content, specifically altering the frequencies for 64C overall and 32C locally in the synaptic myonuclei, yet increased ploidy persists in the synaptic compared to the non-synaptic myonuclei.

**Figure 4.**
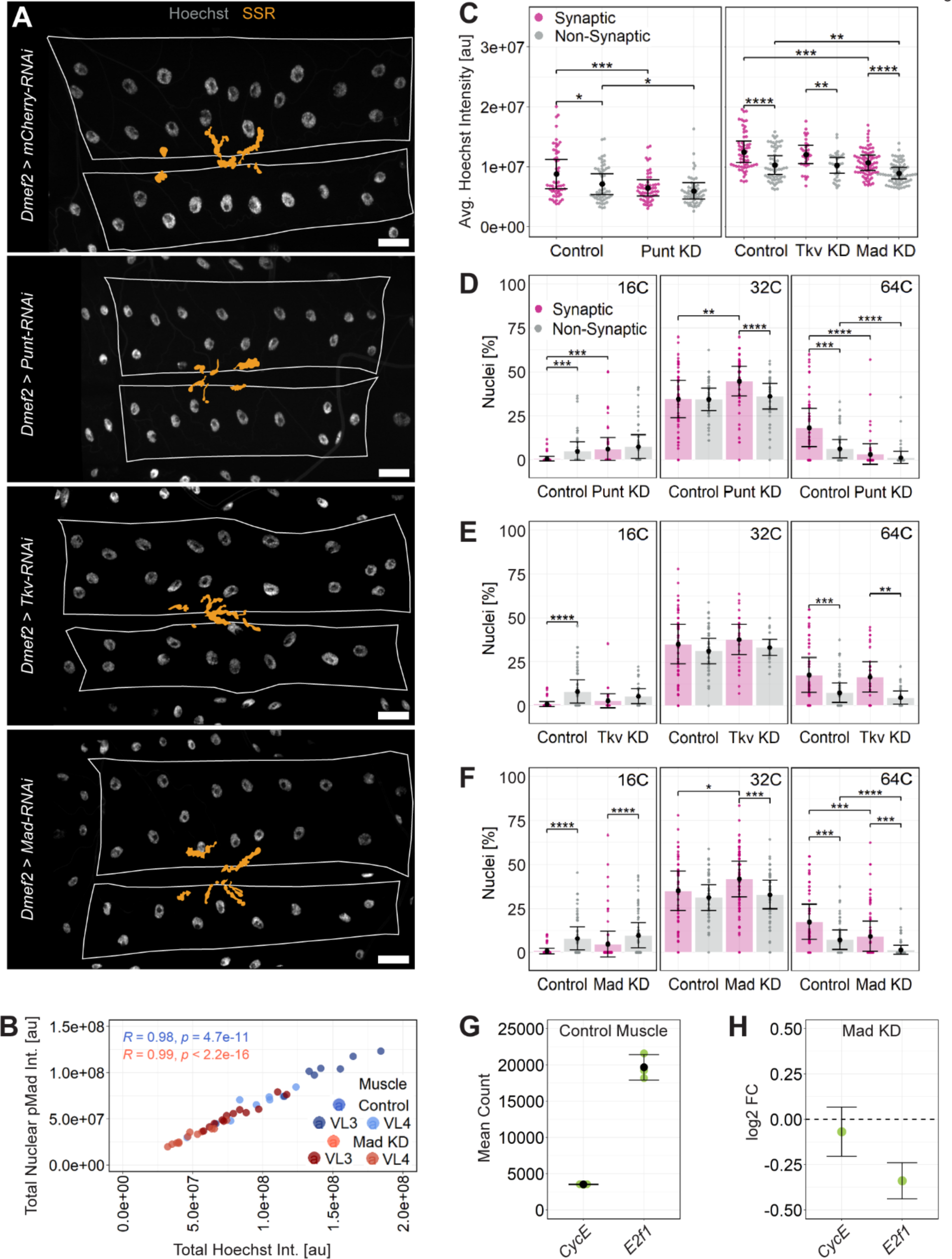
Postsynaptic BMP signaling regulates myonuclear DNA content and endoreplication. **(A)** Confocal images (sum slice projection) of control and Punt, Tkv, and Mad KD (KD) muscles labeled with Hoechst (DNA, white) and anti-Dlg (NMJ, orange mask). Scale bars = 30 µm. **(B)** Total/cumulative nuclear pMad intensity plotted against total/cumulative Hoechst intensity for control and Mad KD VL muscles. Pearson’s correlation coefficient (R) and statistical significance (p) are indicated and reveal a strong linear correlation between nuclear pMad and DNA content for both genotypes (control: *n =* 6 larvae, 16 muscles; Mad KD: *n =* 9 larvae; 24 muscles). **(C)** Average Hoechst intensity per nucleus from control and Punt KD VL3/4 muscles (*n =* 17 larvae, 56 muscles in both groups), and control, Tkv, and Mad KD muscles (control: *n =* 19 larvae, 60 muscles; Tkv KD *n =* 12 larvae, 30 muscles; Mad KD *n =* 20 larvae, 76 muscles). **(D-F)** Frequency of 16C, 32C, and 64C ploidy in synaptic and non-synaptic myonuclei of **(D)** Punt KD, **(E)** Tkv KD, and **(F)** Mad KD VL muscles and corresponding controls. **(G)** Normalized mRNA counts of endoreplication regulators *CycE* and *E2f1* in muscle-enriched preps of control larvae (*N =* 3 replicates, *n =* 6-10 larvae). (H) Fold change of *CycE* and *E2f1* expression in Mad KD preps (*N =* 3 replicates, *n =* 6-10 larvae per genotype). Data shown as mean ± SEM, except in plots G and H where mean ± SD is shown. Statistical comparisons were made for all groups with significant differences noted (* p < 0.05, ** p < 0.01, *** p < 0.001, **** p < 0.0001).

To further explore the role of BMP signaling, we performed bulk RNA sequencing on control and Mad KD muscle-enriched larval preparations. As expected, Mad KD muscles had reduced expression of *Mad* (log2FC = −0.56; *p =* 0.00018) consistent with a ∼30% reduction as found by qPCR. Expression of *brk*, a known Mad target gene that is normally downregulated by Mad (Jaźwińska et al., 1999; Müller et al., 2003; Pyrowolakis et al., 2004) was increased in Mad KD muscles (log2FC = 1.04; *p =* 4.79e-07) (Fig. S3B). Consistent with reduced sizes of Mad KD muscles, GO analysis for biological processes showed downregulation of metabolic pathways (e.g., carbohydrate catabolic process, glucose metabolic process, and ATP generation from ADP) involved in nutrient and cellular energy production that contribute to muscle growth (Demontis and Perrimon, 2009; Piccirillo et al., 2014) (Fig. S3C). Conversely, muscle Mad KD led to an upregulation in genes related to developmental processes (e.g., metamorphosis, post-embryonic organ morphogenesis) and transcription/translation (e.g., cellular response to unfolded protein, polytene chromosome puffing) (Fig. S3D). These data support a growth-promoting role for postsynaptic BMP signaling and suggest that changes in muscle size and myonuclear function may induce a cell stress response. In addition, we examined the expression of muscle-specific structural genes (e.g., *Tm1,* Tropomyosin 1; *Mlc2,* Myosin light chain 2; *Mhc,* Myosin heavy chain) that contribute to muscle mass and observed most of these genes were significantly downregulated in Mad KD muscles (Fig. S3E), further supporting that changes in myonuclear function can affect muscle growth.

To address changes in myonuclear DNA content following Mad KD, we examined regulators of endoreplication. The transcription factor E2f1 is required to drive the expression of cell cycle genes like CycE that trigger S phase initiation. Conversely, E2f1 protein degradation and reduction of CycE activity resets the cell cycle to G phase (Zielke et al., 2011). In control muscles, *E2f1* was highly expressed, whereas *CycE* was moderately expressed (Fig. 4G). Muscle-specific Mad KD significantly reduced *E2f1* expression (log2FC = −0.34; *p =* 0.0157), whereas *CycE* expression was not affected (*p =* 0.9566) (Fig. 4H). qPCR on larval muscle-enriched preps revealed reduced levels of *E2f1* and increased *brk* levels with Mad KD, confirming our RNA sequencing data (Fig. S3F). Further support for this Mad-E2f1 relationship was obtained by analysis of open access databases by JASPAR (2018 CORE collection; Khan et al., 2018) and the Swiss Institute of Bioinformatics (Eukaryotic Promoter Database, EPDnew; Dreos et al., 2015) for *D. melanogaster* that maps transcription factor binding motifs to the upstream promoter and intron regions of genes. Using these databases, we cross-referenced *E2f1* hits for the Mad binding motif and found several potential Mad binding sites within *E2f1* regulatory regions. Thus, it is possible that BMP signaling directly promotes endoreplication by regulating *E2f1* transcription. As a reduction in myonuclear DNA content could also indirectly affect muscle growth and affect the expression of genes required for NMJ structure and function, we next evaluated NMJ gene expression and the synapse itself.

### Postsynaptic BMP signaling shapes NMJ size

Retrograde BMP signaling has been viewed as the main BMP pathway at the NMJ, contributing to NMJ growth and differentiation. Only one study concluded that postsynaptic BMP signaling affects bouton number and thereby NMJ size (Fuentes-Medel et al., 2012). To expand upon this finding, we examined the NMJ size effects of muscle-specific KD of canonical BMP signaling (Tkv, Mad) without affecting the Activin pathway that shares a type II receptor (Punt). To first test whether our manipulations affect the retrograde BMP pathway, we performed immunostaining for BMP ligand Gbb. While Gbb is expressed in both muscles and motor neurons, muscle Gbb is seen as the critical ligand for retrograde pathway activation (McCabe et al., 2003) (Fig. 5A). We found a similar number of Gbb-positive puncta in Tkv and Mad KD muscles compared to control (Fig. 5B). Moreover, by RNA sequencing we observed no change in *gbb* expression in Mad KD muscles (Fig. S4A). These data suggest that Gbb production is unaffected in Tkv and Mad KD muscles, and that Gbb is available to activate the retrograde BMP pathway independent of postsynaptic BMP signaling inhibition.

**Figure 5.**
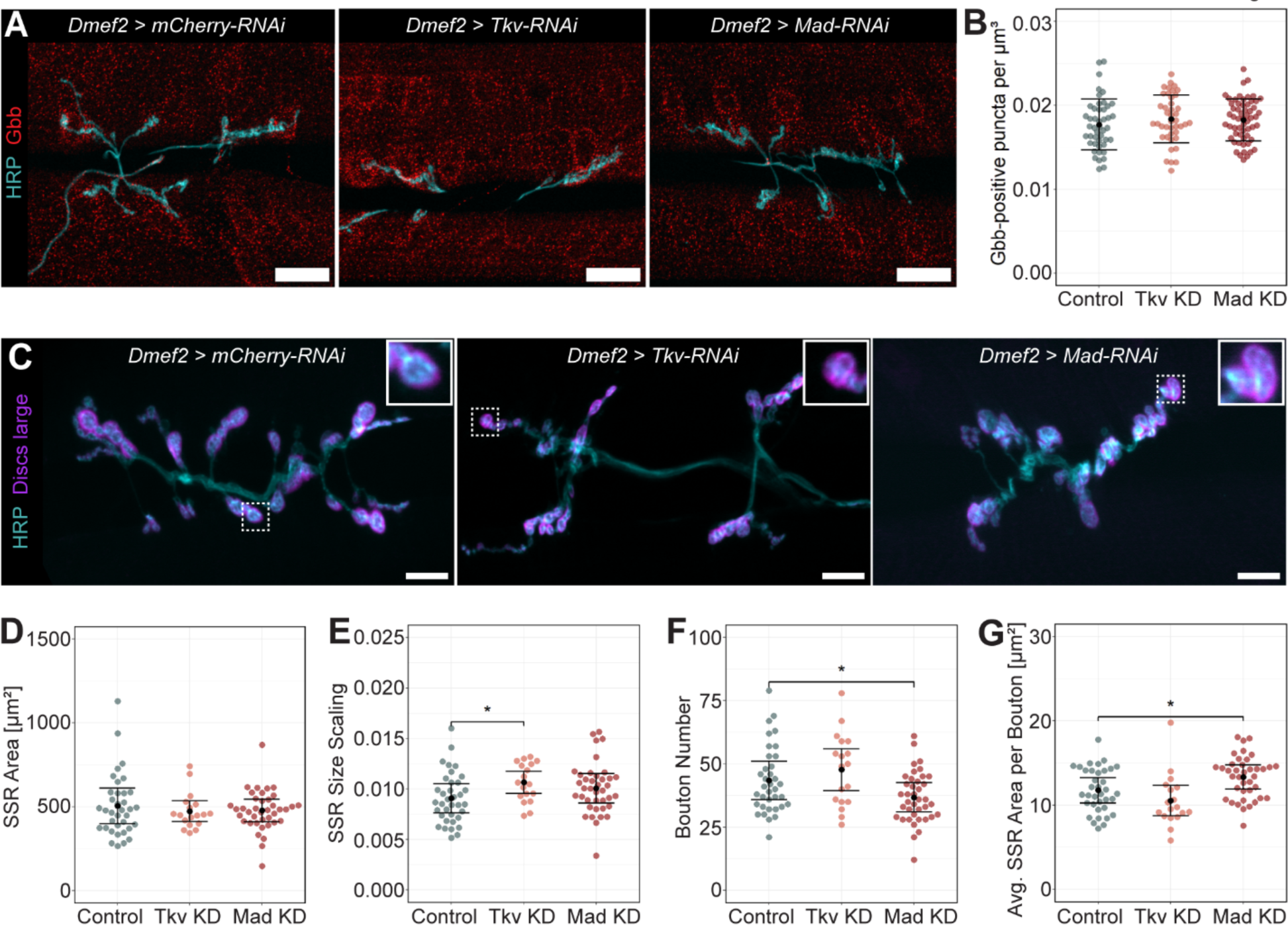
Postsynaptic BMP signaling acts independent of the retrograde BMP pathway to influence NMJ structure. **(A)** Confocal images (maximum intensity projection) showing muscles of the indicated genotypes labeled for presynaptic NMJ marker HRP (cyan) and BMP ligand Gbb (red). Scale bars = 30 µm. **(B)** Quantification of Gbb density (puncta per µm^3^) in VL3 and VL4 muscles of the indicated genotypes (control: *n =* 10 larvae, 46 muscles; Tkv KD: *n =* 12 larvae, 42 muscles; Mad KD: *n =* 13 larvae, 58 muscles). Data shown as mean ± SD. **(C)** Confocal images (sum slice projection) of VL3/VL4 NMJs showing postsynaptic (anti-Dlg, magenta) and presynaptic (anti-HRP, cyan) structures in control, Tkv, and Mad KD muscles. For each genotype, an individual 1b bouton and surrounding SSR (dashed line boxes) is magnified in the top right corner. Scale bars = 10 µm. **(D-G)** Quantification of **(D**) SSR area based on anti-Dlg labeling, **(E)** SSR size scaling (SSR area normalized to total VL3/VL4 muscle cell area), **(F)** 1b bouton number, and **(G)** SSR area per bouton (SSR area divided by 1b bouton number). Plots show mean ± SEM (*n =* 36 control, 18 Tkv KD, and 39 Mad KD NMJs). Statistical comparisons were made for all groups with significant differences noted (* p < 0.05).

To assess postsynaptic changes in Tkv and Mad KD muscles, we measured total SSR area for VL3 and VL4 using Dlg immunostaining (Fig. 5C). We found that SSR area was similar in Tkv KD and Mad KD muscles compared to control (Fig. 5D). Since NMJ size correlates with muscle size (Ho and Treisman, 2020), and KD of BMP signaling components reduces muscle size (Fig. S2D), we normalized total SSR area by cell area. We found proportionally increased SSR area in Tkv KD muscles yet similar proportions in Mad KD muscles (Fig. 5E), indicating a disruption in the coordination of NMJ size with muscle size downstream of postsynaptic BMP signaling.

We next counted the number of large 1b boutons at the VL3/VL4 NMJ based on Dlg labeling and observed reduced bouton number with Mad KD but not Tkv KD (Fig. 5C inserts, Fig. 5F). We divided total SSR area by bouton number to approximate average SSR area per bouton and found that in Tkv KD muscles, SSR area per bouton was unchanged, while Mad KD muscles had increased SSR area per bouton compared to control (Fig. 5G). Together these results suggested that postsynaptic BMP signaling acts independently of the retrograde pathway to coordinate NMJ and muscle size and promote NMJ growth. KD of the upstream Tkv receptor affects muscle size but not bouton number or SSR size, while KD of the downstream effector Mad has additional consequences on DNA ploidy (Fig. 4) and NMJ structure.

### Postsynaptic BMP signaling impacts NMJ function

To explore the role of BMP signaling in the regulation of NMJ function, we first examined the expression of NMJ-related genes in our RNA sequencing dataset. At the glutamatergic *Drosophila* NMJ, ionotropic glutamate receptors (GluRII) in the SSR allow for postsynaptic calcium (Ca^2+^) influx and muscle membrane depolarization following neurotransmitter binding (reviewed in Menon et al., 2013). An individual GluRII consists of subunits C, D, and E, in addition to either A or B that together are required for proper receptor expression and function at the NMJ (Qin et al., 2005). Strikingly, we found changes in the expression of several GluRII subunits with muscle Mad KD (Fig. 6A): the expression of *gluRIIE* (log2FC = −0.6710; *p =* 3.68e-11) and *gluRIIC* (log2FC = −0.5763; *p =* 0.0012) were reduced, *gluRIID* (log2FC = 0.5040; *p =* 0.0001) was increased, and *gluRIIA* and *gluRIIB* were unchanged (Fig. 6A). These RNA sequencing results were tested and confirmed by qPCR (Fig. S4B).

**Figure 6.**
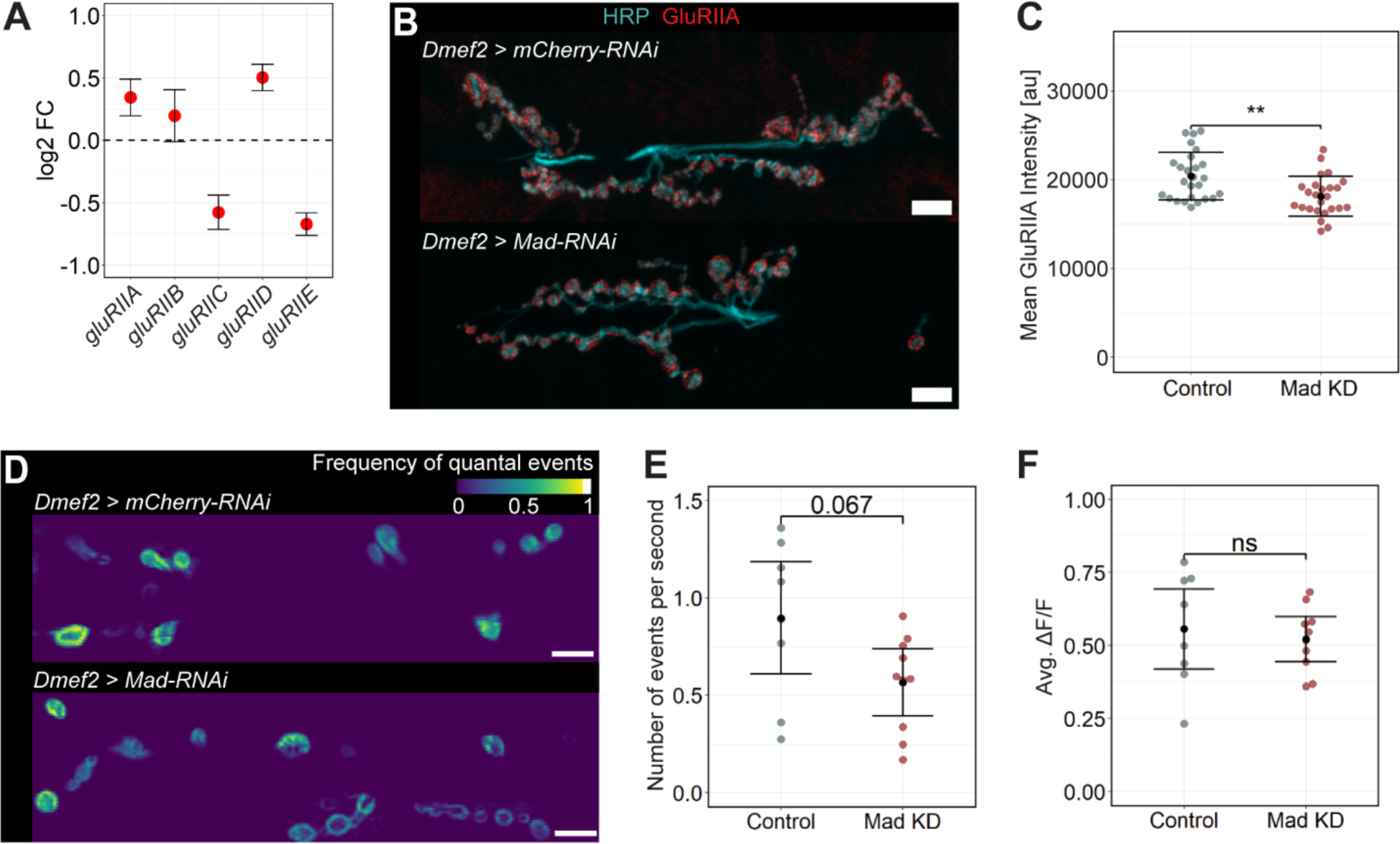
Postsynaptic BMP signaling influences NMJ gene expression and NMJ function. **(A)** Fold change of Glutamate receptor (GluRII) subunits in Mad KD compared to control muscles from RNA sequencing analysis (*N =* 3 replicates, *n =* 6-10 larvae per genotype). Data shown as mean ± SD. **(B)** Confocal images (sum slice projection) of the VL3/VL4 NMJ labeled for HRP (cyan) and GluRII subunit A protein (GluRIIA, red) in control and Mad KD muscles. Scale bars = 10 µm. **(C)** Mean GluRIIA intensity at control and Mad KD NMJs (*n =* 26 control and 25 Mad KD NMJs). Data shown as mean ± SD (** p < 0.01). **(D)** Confocal images (sum slice projection) of control and Mad KD NMJs expressing Ca^2+^ indicator GCaMP6f with SSR localization (*Mhc-CD8-GCaMP6f-Sh*). Pseudo-coloring indicates the frequency of quantal events, defined as the change in GCaMP6f fluorescence from baseline (ΔF/F) above a threshold value, at the NMJs of control and Mad KD muscles over time (20-40 sec). Scale bars = 10 µm. **(E, F)** Quantification of quantal event **(E)** frequency and **(F)** amplitude at the NMJs of control and Mad KD muscles (*n =* 8 control and 10 Mad KD NMJs). Data shown as mean ± SEM.

GluRIIA was previously reported to be markedly reduced at the protein level following muscle-specific Mad KD (Sulkowski et al., 2016). We confirmed these data, finding significantly reduced GluRIIA protein at the NMJs of Mad KD muscles (Fig. 6B-C). In combination with the RNA results, these data suggest that GluRIIA is not directly regulated by Mad transcriptional activity. Regulation of *gluRIIA* expression has been suggested to involve mRNA localization to the SSR and local translation driven by activity of eIF4E, a family of translation initiation factors (Menon et al., 2004). We found a member of the eIF4E family, eIF4E3, was significantly downregulated (log2FC= −6.4810, *p =* 2.95e-15) in Mad KD muscles by RNA sequencing and qPCR (Fig. S4C-D), indicating impaired protein translation at the postsynapse.

The absence or impaired function of GluRIIA inhibits the muscle’s response to spontaneous glutamate release by the motor neuron, resulting in decreased amplitude (quantal size) and frequency of spontaneous miniature excitatory junctional potentials (mEJPs) in the postsynaptic muscle (DiAntonio et al., 1999; Petersen et al., 1997). Single action potentials as well as spontaneous release of neurotransmitter from the motor neuron result in Ca^2+^ influx at the postsynaptic membrane and can be measured using Ca^2+^ indicator GCaMP6f localized to the SSR (*Mhc-CD8-GCaMP6f-Sh*) (Desai and Lnenicka, 2011; Newman et al., 2017). To determine if neurotransmission is affected in Mad KD muscles, we performed live imaging of postsynaptic Ca^2+^ influx during spontaneous mEJPs (Fig. 6D). We found a statistical trend for reduced frequency of quantal events in Mad KD muscles compared to control (Fig. 6E), whereas the amplitude was unchanged (Fig. 6F). This mild functional phenotype is consistent with a reduction but not a loss in GluRIIA protein at the postsynapse and suggests that BMP signaling promotes the postsynaptic response to NMJ neurotransmission.

## Discussion

Each skeletal myofiber in mammals contains hundreds of myonuclei, all sharing one cytoplasm and contributing to optimal muscle function. Several myonuclei are physically clustered at the neuromuscular junction (NMJ) and have a distinct transcriptional profile. Whether these myonuclear specializations are conserved in *Drosophila* and how synaptic myonuclei coordinate their gene expression with NMJ requirements is unclear. In this study, we identified synaptic myonuclei in *Drosophila* larval muscles based on proximity to the NMJ, increased nuclear size scaling, and increased DNA content. Postsynaptic BMP signaling results in an intracellular gradient of nuclear pMad with the highest signal in synaptic myonuclei, which regulates endoreplication, via regulation of *E2f1,* and NMJ-related genes (i.e., GluRII subunits). Our data suggest that local activation and resulting BMP signaling events affect muscle cell size, nuclear size, DNA content, and synaptic structure/size and activity. Altogether, this study reveals how local signaling in a syncytial cell orchestrates nuclear activity on different levels for optimal cell function.

### Synaptic myonuclei in *Drosophila* larval muscles

The skeletal muscle’s postsynapse is characterized by the accumulation of membrane and proteins necessary for synaptic integrity and signal transmission. Mammalian data suggest that growth and maintenance of the postsynaptic protein milieu requires local production of specific mRNAs that are provided by (sub)synaptic myonuclei (Belotti and Schaeffer, 2020). As a result, synaptic myonuclei are important contributors to NMJ activity. Here we identify synaptic myonuclei at the *Drosophila* NMJ by their distance to the NMJ (Fig. 1, S1), like previous studies in mammalian myofibers (Bai et al., 2022; Grady et al., 2005). We uncover several specific adaptations in *Drosophila* synaptic myonuclei: (1) increased nuclear size scaling, signifying that each myonucleus has ample potential to meet local signaling and metabolic demands; (2) Greater DNA content, which increases transcriptional capacity and is correlated with increased H3K9ac, a marker of gene activation, and nucleolar size, a marker of translational potential (Windner et al., 2019); and (3) Higher levels of BMP signaling (pMad) indicating differential transcription in synaptic compared to non-synaptic myonuclei.

In contrast to mammals, *Drosophila* synaptic and non-synaptic myonuclei have similar internuclear distances and do not cluster under control conditions. However, a previous study reported specific nuclear spacing for NMJ-adjacent myonuclei in VL3 muscles, that is regulated by microtubules and associated motor proteins (Perillo and Folker, 2018). We observed a VL3-specific increase in the percent of synaptic nuclei for Tkv KD and Mad KD muscles, suggesting that *Drosophila* muscles use nuclear positioning to compensate for defective synaptic nuclear properties. Altogether, our data strongly suggest that myonuclear changes at the NMJ are a conserved muscle feature. While mammalian synaptic myonuclei are visibly clustered (Hansson et al., 2020), *Drosophila* larval muscles establish a gradient of nuclear size scaling and DNA content to achieve local nuclear adaptations.

A recent study in mammals identified a reliable protein marker for synaptic myonuclei, the nuclear anchoring protein Nesprin-1, that when used in combination with proximity to the NMJ can better distinguish synaptic from non-synaptic myonuclei than NMJ proximity alone (Ruiz et al., 2023). In *Drosophila*, no protein has yet been uncovered to distinguish synaptic myonuclei. While our study indicates pMad as a promising marker, a fixed structural protein would be better than a diffusible signaling effector to identify these nuclei. In the absence of such a protein marker, it is possible that we are overestimating the number of synaptic myonuclei per NMJ or masking subpopulations within the synaptic myonuclei. Nuclei that spatially overlap with the SSR could experience shorter-range NMJ signals that may have additional effects on gene expression. Future work identifying markers specific to synaptic myonuclei in *Drosophila* would be highly useful to corroborate our distance-based identification approach. Importantly, our work indicates several measures (distance, morphology, and DNA ploidy) to identify and study synaptic myonuclei in *Drosophila*, thereby providing an important platform for future studies.

### BMP signaling creates an intracellular pMad gradient

In mammals, synaptic myonuclei transcribe genes encoding NMJ components in response to local signals which relay information on synaptic activity (DeChiara et al., 1996; Ruegg and Bixby, 1998; Sanes and Lichtman, 2001). Of the many local signals, we investigated the BMP pathway, as it signals with the mammalian postsynaptic regulator MuSK to promote BMP target gene expression (Jaime et al., 2024; Yilmaz et al., 2016) and to regulate overall NMJ structure and postsynaptic localization of voltage-gated sodium channels (Nav1.4) required for muscle fiber excitability (Fish et al., 2023). As in mammals, BMP receptors are also found at the postsynaptic membrane of *Drosophila* muscles (Chou et al., 2013; Dudu et al., 2006). We measured BMP signaling activity by quantifying nuclear pMad, the active BMP transcriptional effector, and observed a striking intracellular gradient, with the highest levels within the synaptic myonuclei and the lowest levels within the non-synaptic myonuclei (Fig. 2).

While signaling gradients are usually established extracellularly by the distribution of ligands to transmit positional information, nuclear protein gradients have been described for transcription factors, specifically Bicoid, in the *Drosophila* syncytial blastoderm (Ali-Murthy and Kornberg, 2016). In muscle, the robustness of the pMad gradient is remarkable given that muscle cells continuously contract, which likely results in cytoplasm mixing (Koslover et al., 2017). A similar phenomenon occurs in mammalian myofibers for transcription factor GABP that is globally expressed but specifically phosphorylated by local signaling pathways to activate NMJ gene transcription in synaptic myonuclei (Belotti and Schaeffer, 2020). By establishing a transcription factor gradient, *Drosophila* synaptic myonuclei would achieve higher transcription of pMad target genes than non-synaptic myonuclei. These transcriptional differences could then locally promote adaptions that affect synaptic myonuclear output and NMJ functionality.

### BMP signaling affects myonuclear size and DNA content

As synaptic myonuclei show an enrichment for pMad, we dissected the postsynaptic BMP signaling pathway to understand the role of BMP signaling in synaptic myonuclear adaptations.

Muscle KD of each BMP pathway component (Punt, Tkv, Mad) resulted in smaller muscles (i.e., smaller cytoplasmic domain sizes) and smaller nuclei, but maintained most differences between synaptic and non-synaptic nuclei (Fig. 3-4, S2). In addition, we found genotype-specific phenotypes that further illuminate BMP signaling effects. Overall, KD of Tkv had no or mild effects on size and DNA content/ploidy and did not alter differences between synaptic and non-synaptic nuclei. This could be attributed to a smaller reduction in BMP signaling due to several factors: (1) achieving only ∼20% KD of highly expressed Tkv in larval muscles to allow for sufficient Tkv activity and/or (2) larval muscles expressing two type I BMP receptors, Tkv and Sax, thereby Sax-mediated signaling may provide some compensation.

Punt KD in muscles resulted in a robust reduction in size and DNA content (less 64C, more 32C ploidy); however, the effects on DNA content were stronger in synaptic than non-synaptic nuclei. Punt is believed to be the only BMP type II receptor expressed in larval muscle (Akiyama et al., 2024), where it functions to stabilize ligand binding and phosphorylate the type I receptor for activation. However, Punt also associates with Activin receptors (Babo), and like BMP, this other TGF-β pathway has been shown to influence NMJ size and activity (Ellis et al., 2010; Fuentes-Medel et al., 2012; Kim and O’Connor, 2014). It will be of interest in future studies to assess whether Activin signaling through Punt-Babo receptor activity regulates intracellular heterogeneity by specifically affecting synaptic myonuclear adaptions.

Muscle KD of Mad, the transcriptional effector of BMP signaling, reduced nuclear pMad levels by 50% and showed the most robust effects on size and DNA content with significant DNA reduction in synaptic and non-synaptic nuclei. In addition to overall reduction in muscle size and nuclear size, Mad KD results in reduced nuclear size scaling specifically in the synaptic myonuclei.

To provide sufficient gene products to the cytoplasmic domains as well as the NMJ, transcription and/or translation would need to be upregulated. Indeed, we observe that pathways related to transcription and translation are upregulated (e.g., cellular response to unfolded protein, polytene chromosome puffing) in Mad KD muscles. However, expression of *eIF4E3*, a component of the postsynaptic translational machinery localized to the NMJ, and a locally translated protein, GluRIIA subunit, are significantly reduced in Mad KD muscles. Therefore, these data suggest insufficient compensation of synaptic nuclei in Mad KD muscles to meet the demand for critical postsynaptic proteins.

Our data indicate that BMP signaling directly regulates DNA endoreplication and thus nuclear DNA content. In support, we found reduced expression of the endoreplication regulator *E2f1* with muscle Mad KD, and potential Mad binding sites in *E2f1* regulatory regions. E2f1 promotes the G1-S phase transition and is required for DNA replication in cells undergoing mitosis and endoreplication (Duronio et al., 1995). *E2f1* transcription is not under cell cycle control, but levels of E2f1 protein and its regulated transcripts (e.g., *CycE*) do oscillate with the cell cycle due to post-transcriptional regulation of E2f1 (Øvrebø et al., 2022; Zielke et al., 2011). This specific translational control of E2f1 function may explain why our BMP manipulations only impaired but did not disrupt endocycling. Like in flies, myonuclei in mammals are recently understood to undergo endoreplication to increase DNA content (Borowik et al., 2022), but the regulation of this process is unknown. BMP regulation of E2f1 expression may be generalizable to muscle and other polyploid cells since BMP signaling was recently shown to intersect with E2f1 activity to promote endoreplication in *Drosophila* prostate-like secondary cells (Sekar et al., 2023). In larval muscle, the pMad gradient correlates with ploidy distribution within the cell, thus differing amounts of BMP target gene expression and endoreplication could be involved in establishing myonuclear heterogeneity.

A deficit in nuclear DNA content could indirectly affect nuclear size and muscle growth; however, we also observed downregulation of multiple structural genes in Mad KD muscles that encode components of the myofiber’s contractile apparatus (Fig. S3). Of note, E2f1 was shown in *Drosophila* adult skeletal muscle to regulate muscle mass by controlling the expression of several of these structural genes (e.g., *Tm1*) (Zappia and Frolov, 2016). Furthermore, we observed a downregulation of genes involved in metabolic pathways in Mad KD muscles that are known to promote muscle growth (Demontis and Perrimon, 2009; Piccirillo et al., 2014). Thus, BMP signaling could affect muscle cell size directly through promoting nuclear expression of structural and metabolic genes and/or indirectly via E2f1 and regulation of DNA content.

### Postsynaptic BMP signaling shapes synaptic myonuclear contributions to NMJ size and function

BMP signaling at the *Drosophila* NMJ is largely studied as a retrograde pathway that activates presynaptic BMP receptors and drives neuronal gene expression to promote NMJ growth through modulating presynaptic actin dynamics and bouton budding (Aberle et al., 2002; Ball et al., 2010; Marqués et al., 2002; McCabe et al., 2003; Piccioli and Littleton, 2014). Our findings suggest that postsynaptic BMP signaling is independently playing a role in NMJ size regulation. We find that KD of Tkv disrupts the size relationship between the muscle and the NMJ/SSR. This phenotype does not occur in Mad KD muscles and may be driven by non-canonical Tkv activity. Non-canonical BMP signaling at the NMJ has previously been reported for the neuronal type II receptor Wit that can associate with actin regulator Limk to regulate the NMJ (Piccioli and Littleton, 2014); therefore, it is possible that non-canonical BMP pathways also exist postsynaptically in the muscle. Mad KD led to a reduction in the number of boutons and greater SSR area per bouton (Fig. 5). An increase in SSR area could result from the formation of satellite boutons, which are small ectopic boutons with a common surrounding SSR that develop from the axon or a single bouton (Guangming et al., 2020). Satellite boutons develop in various endocytic mutants, which are known to alter NMJ neurotransmission (Dickman et al., 2006). However, our results are based on confocal microscopy and further ultrastructural studies will be needed to confirm changes in presynaptic bouton structure in Mad KD muscles.

Our data indicate that postsynaptic BMP signaling also affects the expression of Glutamate receptor (GluRII) subunits at the mRNA and protein level, as well as the frequency of spontaneous quantal events, signifying an impaired postsynaptic response to neurotransmitter release (Fig. 6). This is consistent with previous work suggesting that postsynaptic BMP signaling impacts neurotransmission at the *Drosophila* NMJ through altering GluRII protein levels (Rawson et al., 2003; Sulkowski et al., 2016). Further, loss of GluRIIA (*GluRIIA^SP16^)* results in a reduced postsynaptic response to spontaneous neurotransmitter release, measured by a lower amplitude (quantal size) and a lower frequency of mEJPs (Petersen et al., 1997). We suggest that postsynaptic BMP signaling supports proper NMJ activity through affecting GluRII subunit protein levels both through transcriptional regulation by Mad and by promoting the expression of postsynaptic translational machinery (i.e., eIF4E3) involved in local GluRII subunit protein translation.

Altogether, our study works toward a more comprehensive understanding of BMP pathway activation at the NMJ and introduces muscle-specific adaptions in *Drosophila* that include changes in DNA content and size scaling of postsynaptic myonuclei. We propose that local postsynaptic BMP signaling provides input on NMJ size and function through affecting endoreplication and/or the expression of genes involved in NMJ function. In mammals, loss of synaptic myonuclei occurs with aging and corresponds with NMJ structural and functional defects. A recent study revealed that increasing synaptic myonuclear number by muscle activation of 4EBP1, a mRNA translation regulator, led to increased acetylcholine receptor subunit expression and neurotransmission in aged mice (Ang et al., 2022). Postsynaptic BMP signaling could also be a good target in sarcopenia, an aging-related process of muscle deterioration and other conditions that implicate NMJ activity in their initiation and/or disease progression.

## Materials and methods

### *Drosophila* husbandry, stocks, and crosses

Experimental crosses of *Drosophila melanogaster* were grown on standard cornmeal medium at 25 °C in 12-hour light/12-hour dark conditions under humidity control. *Drosophila* stocks were maintained at room temperature. RNA interference and overexpression studies relied on the Gal4-UAS system (Brand and Perrimon, 1993). For muscle-specific expression, we used the *Dmef2-Gal4* driver line paired with the following *UAS* lines that target core components of the BMP signaling pathway: *UAS-Mad-RNAi* [Vienna Drosophila Resource Center (VDRC) 12635], *UAS-tkv-RNAi* [Bloomington Drosophila Stock Center (BDSC) 35653], and *UAS-put-RNAi* [VDRC 848]. The driver lines were crossed to *UAS-mCherry-RNAi* [BDSC 35785] as the control RNAi. We recombined the *Dmef2-Gal4* driver line with *Mhc-CD8-GCaMP6f-Sh* line (Newman et al., 2017) [BDSC 67739] to create *Dmef2-Gal4, Mhc-CD8-GCaMP6f-Sh* flies for quantal live imaging.

### Larval staging and muscle selection

For all experiments, larvae were raised at 25 °C in standard food vials, and both male and female larvae were used at the wandering third instar stage. Staging was confirmed using the developmental landmarks of mouth hook and spiracle morphologies (Bodenstein, 1950). Larval body wall muscles ventral longitudinal 3 and 4 (VL3 and VL4) were studied in anterior abdominal hemi-segments 2 and 3 due to having a consistent muscle size and NMJ span and size.

### RNA isolation and qRT-PCR

Six to fifteen larvae at the third instar wandering stage were dissected and rinsed in ice-cold HL3.1 medium (Feng et al., 2004) with all internal organs removed and the larval body wall kept intact as previously described (Brent et al., 2009). The head and tail portions of the larva were also removed to enrich for muscles in the body wall. Total RNA was isolated from muscle-enriched larval fillets using TRIzol reagent (Thermo Fisher Scientific, 15596026) and chloroform extraction, followed by purification with TURBO DNA-free Kit (Thermo Fisher Scientific, AM1907). cDNA was synthesized from 1 µg of RNA using the SuperScript III First-Strand Synthesis System for RT-PCR (Thermo Fisher Scientific, 18080-051), and qRT-PCR reactions performed using the CFX96 Real-Time PCR system (BioRad) with SYBR Select Master Mix for CFX (Applied Biosystems, 4472942). Three independent mRNA preparations per genotype were made and plated in triplicate. Relative gene expression was calculated relative to *RpL32* by the delta-delta Ct method (Livak and Schmittgen, 2001; Schmittgen and Livak, 2008). Data are shown as fold change in gene expression and ΔCt values were used for statistical comparison. Table S1 contains information on the oligonucleotides used.

### RNA sequencing and analysis

Six to ten wandering third instar larvae per genotype were dissected as described above for each replicate of three in total. Total RNA was isolated using TRIzol and chloroform extraction from the muscle-enriched larval fillets. The lysates were submitted to the Memorial Sloan Kettering Cancer Center (MSKCC) Integrated Genomics Operations (IGO) core for RNA quality assessment and Illumina next-generation sequencing. The sequencing libraries were constructed using Poly A enrichment following oligo-dT-mediated purification. The libraries were then sequenced on the Illumina HiSeq 4000 in a PE50 run. Over 40 million reads were generated per library. FASTQ data was processed in R with the ShortRead package (Morgan et al., 2009) and quality control was conducted with the Rfastp package (Wang and Carroll, 2023). Reads were mapped and aligned to the *Drosophila* genome (release dm6) using Rsubread (Liao et al., 2019) and counted with the GenomicAlignments package (Lawrence et al., 2013). Gene differential expression analysis was performed using DESeq2 (Love et al., 2014). Lastly, enrichment analysis was performed with GOseq (Young et al., 2010) using the GO database.

### Immunostaining and confocal imaging

Larvae were dissected to optimally visualize the body wall muscles in ice-cold HL3.1 buffer and fixed with 4% paraformaldehyde in HL3.1 for 20 minutes for all antibodies except GluRIIA for which samples were fixed in Bouin’s fixative (Electron Microscopy Sciences, 15990) for 5 minutes. Larval fillets were blocked in PBT-BSA [0.1% BSA, 0.3% Triton X-100 in PBS pH 7.4] for 30 minutes and incubated with the following primary antibodies overnight at 4 °C: mouse anti-Lamin C (DSHB, LC28.26; 1:50), mouse anti-Discs large (DSHB, 4F3; 1:200-1:300), guinea pig anti-phosphorylated Smad 1/5/8 (a gift from E. Laufer; 1:500; previously shown to be specific to *Drosophila* phosphorylated Mad in Guo et al., 2013), mouse anti-GluRIIA (DSHB, 8B4D2 (MH2B); 1:50), or mouse anti-Gbb (DSHB, 3D6-24; 1:50). Then larval fillets were washed in PBT-BSA and incubated with Alexa Fluor-conjugated secondary antibodies, F-actin probe 488-conjugated Phalloidin, 647-conjugated Horseradish Peroxidase (HRP) that stains insect neuronal membranes, and/or Hoechst 33342 at a 1:400 concentration for 1 hour at room temperature. Lastly, larval fillets were washed in PBT and mounted on slides and cured over 24 hours in ProLong Gold (Invitrogen, P36930). All antibodies used are described in Table S1. Confocal z-stacks of VL3 and VL4 muscles were acquired using an A1R laser-scanning microscope (Nikon) with an oil-immersion objective (CFI Plan APO Lambda S 40x Sil/1.25). The same confocal laser and system settings were used for all samples under direct comparison. Approximately seven larvae of each genotype were dissected, stained, and imaged for each experiment that was repeated for a total of three independent experiments, unless otherwise noted.

### Muscle nuclear size, DNA content, pMad, and position analysis

Sum slice projections of confocal z-stacks were made in ImageJ (FIJI) for 2D quantification of VL3 and VL4 muscles. Muscle areas for VL3 and VL4 were traced by hand as determined by phalloidin labeling using the polygon selection tool. These cell outlines were used to measure muscle cell area and *x*- and *y*-coordinates of the anterior and posterior ends of individual cells. Automated thresholding of fluorescence intensities using Triangle mode of Lamin C and/or Hoechst labeling was used to create binary images of VL nuclei. The number, size (areas), and position (centroids) of all nuclei within each muscle cell were recorded. The binary images served as masks to measure Hoechst and nuclear pMad fluorescence intensities (sum of pixel values in a nuclear area). Hoechst values were used to estimate DNA content and ploidy as we and other labs have previously done (Dej and Spradling, 1999; Losick et al., 2016; Sher et al., 2013; Unhavaithaya and Orr-Weaver, 2012; Windner et al., 2019). In addition, nuclear centroids were used to determine nearest neighbor distances and make Voronoi tessellations to measure cytoplasmic domain areas (Du et al., 2010). Average nuclear values for all nuclei in an individual muscle cell from 6 to 13 muscles per experiment across three replicates were graphed and used for statistical analyses.

### Nuclear distance to the neuromuscular junction (NMJ) measurement

Sum slice projections used for muscle nuclear analyses were also used to generate binary images of the NMJ for the VL3 and VL4 muscles. This was done by automated thresholding of discs large labeling using Triangle mode in FIJI that included both 1b and 1s synapse elements. The shortest distance from multiple objects (nuclear centroids) to one feature (the NMJ) was determined using a FIJI macro developed by Michael Cammer and others at NYU Langone Medical Center (available at https://microscopynotes.com/imagej/shortest_distance_to_line/index.html). The NMJ was divided for VL3 and VL4 based on discs large and phalloidin labeling, and then the shortest nuclear distance to the NMJ was measured for nuclei of an individual muscle cell, calculated separately for VL3 and VL4.

### NMJ size analyses

Sum slice projections of confocal z-stacks of discs large immunostaining were made in ImageJ (FIJI) for 2D quantification of the NMJ for VL3/VL4 muscle pair. We performed automated thresholding of Discs Large fluorescence intensity using Yen mode to create binary images of the VL3/4 NMJ. This thresholding approach produced a binary of 1b synapse components only, purposefully excluding 1s synapse components due to the difficulty of reliably measuring 1s synapse area and number for the VL3/4 NMJ. The size (area) of the discs large-positive subsynaptic reticulum (SSR), the NMJ’s postsynaptic element, was recorded. The number of 1b boutons was quantified by using the counter tool and counting the number of discs large-positive rings that surround 1b boutons, optical slice by optical slice in the confocal z-stack. The average SSR area per bouton was calculated by dividing SSR area by 1b bouton number. The SSR scaling was determined by dividing SSR area by total muscle area for VL3 and VL4, with area determined for the individual muscles as described above (see ‘Muscle nuclear size, DNA content, pMad, and position analysis’). Values for individual VL3/VL4 NMJs (7-13 NMJs per genotype from one experiment) across three replicates were graphed and used for statistical analyses.

### Quantification of Muscle Gbb

In Imaris 10.0 (Bitplane), confocal z-stacks of phalloidin, Gbb, and HRP immunostainings were processed. Surfaces were created manually for VL3 and VL4 muscles separately based on phalloidin labeling. The spots function was used to detect Gbb-positive puncta (diameter equals 1 µm) in 3D under automated thresholding within the VL3 and VL4 surfaces. The number of Gbb-positive puncta and the surface volumes (µm^3^) were recorded for each muscle. The number of Gbb-positive puncta per µm^3^ was calculated by dividing the number of puncta by surface volume for an individual muscle cell. Muscles were excluded from analysis if there was HRP-positive motor axons overlayed on the muscle, an artifact of dissection, since Gbb is expressed both in the muscle and motor neuron and would impact Gbb count. This analysis was conducted in duplicate with 42-58 muscles from 10-13 larvae per genotype.

### Glutamate receptor intensity analyses

In Imaris 10.0 (Bitplane), confocal z-stacks of GluRIIA and HRP immunostainings were processed as previously done (Christophers et al., 2023). A three-dimensional surface for HRP was generated with a surface detail grain level of 0.355 µm, smoothing enabled, and auto-thresholding. Small surfaces were removed that were outside of the NMJ. A mask was created from the HRP surface using the Distance Transform setting, and then this mask was used to create a second surface, with a surface detail level of 0.355 µm and a manual threshold of 0 to 0.5 µm to limit the final surface to a shell extending from the edge of the HRP surface to 0.5 µm away. The sum of the sum intensity of the GluRIIA channel within this 0.5 µm-wide surface was recorded and normalized to the expanded HRP volume. This analysis was conducted in duplicate with 25-26 NMJs from 9 larvae per genotype.

### Quantal live imaging and analysis

Seven to nine wandering third instar larvae were dissected and pinned to visualize the body wall muscles in ice-cold HL3.1 buffer on 5.5-cm wide, sylgard plates. Live VL3/VL4 NMJs in abdominal hemi-segment 3 were imaged in ambient temperature using a Stellaris 5 laser-scanning confocal microscope (Leica) with a water-immersion objective (HC FLUOTAR L VISIR 25x/0.95) and HyD S detector in counting mode. Confocal images of SSR-localized GCaMP6f signal were acquired continuously for two-20 second sessions by scanning bidirectionally at 1000 Hz, with a pixel size per voxel size of 0.728 µm and an area size of 512 x 100 pixels to provide a frame rate of 18.18 frames per second with 2.25x zoom applied in Leica LASX software. The pinhole size was 65.1 µm, calculated at 1.5 A.U. for 451 nm emission. Images were analyzed in FIJI based on a published protocol (Chen et al., 2024) followed by manual setting of the threshold (ΔF/F = 0.25) to separate signal to noise for peak event identification in Python using the sci-kit image package. Quantal event frequency was calculated as the number of peak events over time, and quantal event amplitude was the average fluorescence (ΔF/F) of the peak events. Due to larval movement, only positionally stable NMJs were included for analysis that were collected over a minimum of 20 seconds, ranging from a total of 20 to 40 seconds. This analysis was conducted in duplicate with 8-10 NMJs from 7-9 larvae per genotype.

### Statistical analysis

Correlation coefficients (R), pairwise comparisons between two groups determined by two-tailed student’s *t*-test (alpha of 0.05), and three or more group comparisons determined by one-way ANOVA were computed using R statistical software (version 1.4.1717). Data are shown as mean ± s.e.m., unless otherwise noted, with asterisks used to denote significance level (*p =* * < 0.05, ** < 0.01, *** < 0.001, **** < 0.0001) and the sample size reported in the figure legend. All plots were made using R with the ggplot2 package (Wickman, 2016).

## Supplemental material

Further details about nuclear and RNA analyses in *Drosophila* body wall muscles in control and following muscle-specific BMP manipulations are available in the supplemental material.

## Data availability statement

The data underlying Figures 1-6 and supplemental Figures S1-4 are available in the published article and its online supplemental material. The RNA sequencing data underlying Figures 4 and 6 and supplemental Figures S3-4 are openly available in Gene Expression Omnibus (GEO) [GSE263700].

## Acknowledgements

We thank the Baylies lab members for helpful discussions on data analysis, data interpretation, and giving feedback on the manuscript, V. Basu for coding assistance, and E. Laufer for providing the phosphorylated Mad antibody. We thank the members of the Integrated Genomics Operations Core at MSKCC for their important contributions to this work, the Bloomington and Vienna Drosophila Stock Centers and Zurich FlyORF for genetic fly lines, and the Developmental Studies Hybridoma Bank for antibodies. This work was supported by NIH grant R35GM141877 (M.K.B) and NaÄonal Cancer InsÄtute P30 CA 008748 to MSKCC. V.E.V. was supported by a Medical Scientist Training Program grant from the National Institute of General Medical Sciences of the NIH under award T32GM007739 to the Weill Cornell/Rockefeller/Sloan Kettering Tri-Institutional MD-PhD Program. The authors declare no competing or financial interests.

## Author contributions

1. V. von Saucken conceptualized, performed, and formally analyzed experiments and wrote the original draft and reviewed and edited the final manuscript. S. Windner contributed to experiment methodology and reviewed and edited the final manuscript. M. Baylies supervised the work, acquired funding, and reviewed and edited the final manuscript.

## Abbreviations

BDSC, Bloomington Drosophila Stock Center; BMP, bone morphogenetic protein; brk, brinker; C, copies; Ca^2+^, calcium; CycE, cyclin E; Dlg, discs large; DSHB, Developmental Studies Hybridoma Bank; E2f1, E2F transcription factor 1; eIF4E3, eukaryotic translation initiation factor 4E3; FC, fold change; Gbb, glass bottom boat; GluR, glutamate receptor; KD, knockdown; mEJP, miniature excitatory junctional potential; NMJ, neuromuscular junction; pMad, phosphorylated Mad; Put, Punt receptor; RNAi, RNA interference; Sax, Saxophone receptor; SSR, subsynaptic reticulum; Tkv, Thickveins receptor; VL, ventral longitudinal

---

**Figure S1.**
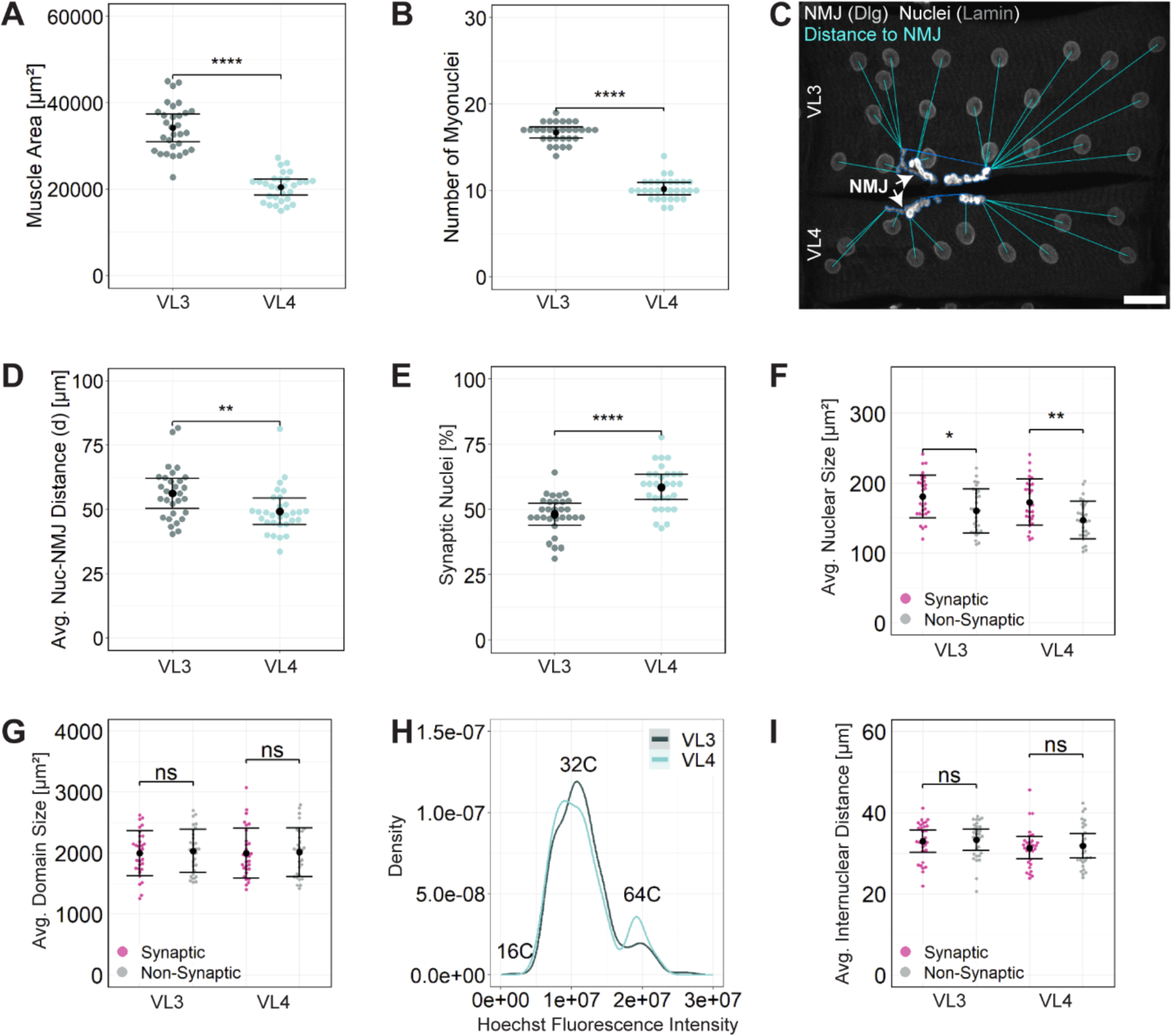
Characterization of synaptic myonuclei in *Drosophila* larval VL3 and VL4 muscles. **(A)** Average VL3 and VL4 muscle area based on actin (phalloidin). **(B)** Myonuclear number for individual VL3 and VL4 muscles. **(C)** Confocal image (sum slice projection) of a VL3/VL4 muscle pair labeled for NMJ marker Dlg (white, arrows) and nuclear envelope protein Lamin C (gray). Cyan lines indicate direct distance between individual myonuclear centers and the nearest NMJ component in VL3 and VL4 muscles. Scale bar = 30 µm. **(D)** Average nuclear distance to the NMJ (d) for VL3 and VL4 muscles. **(E)** Average percent of synaptic myonuclei in VL3 and VL4 muscles. **(F)** Average nuclear size for synaptic and non-synaptic myonuclei in VL3 and VL4 muscles. **(G)** Average cytoplasmic domain size determined by Voronoi tessellations for synaptic and non-synaptic myonuclei from VL3 and VL4 muscles. **(H)** A smooth density estimate plot of Hoechst intensities for VL3 and VL4 myonuclei where corresponding ploidies are denoted (16C, 32C, and 64C). **(I)** Average internuclear distance, determined by measuring the nearest neighbor distance for each myonucleus, for synaptic and non-synaptic myonuclei from VL3 and VL4 muscles. Data shown as mean ± SEM (*n =* 19 larvae, 30 muscle pairs of VL3 and VL4 from abdominal hemi-segments 2 and 3; * p < 0.05, ** p < 0.01, **** p < 0.0001).

**Figure S2.**
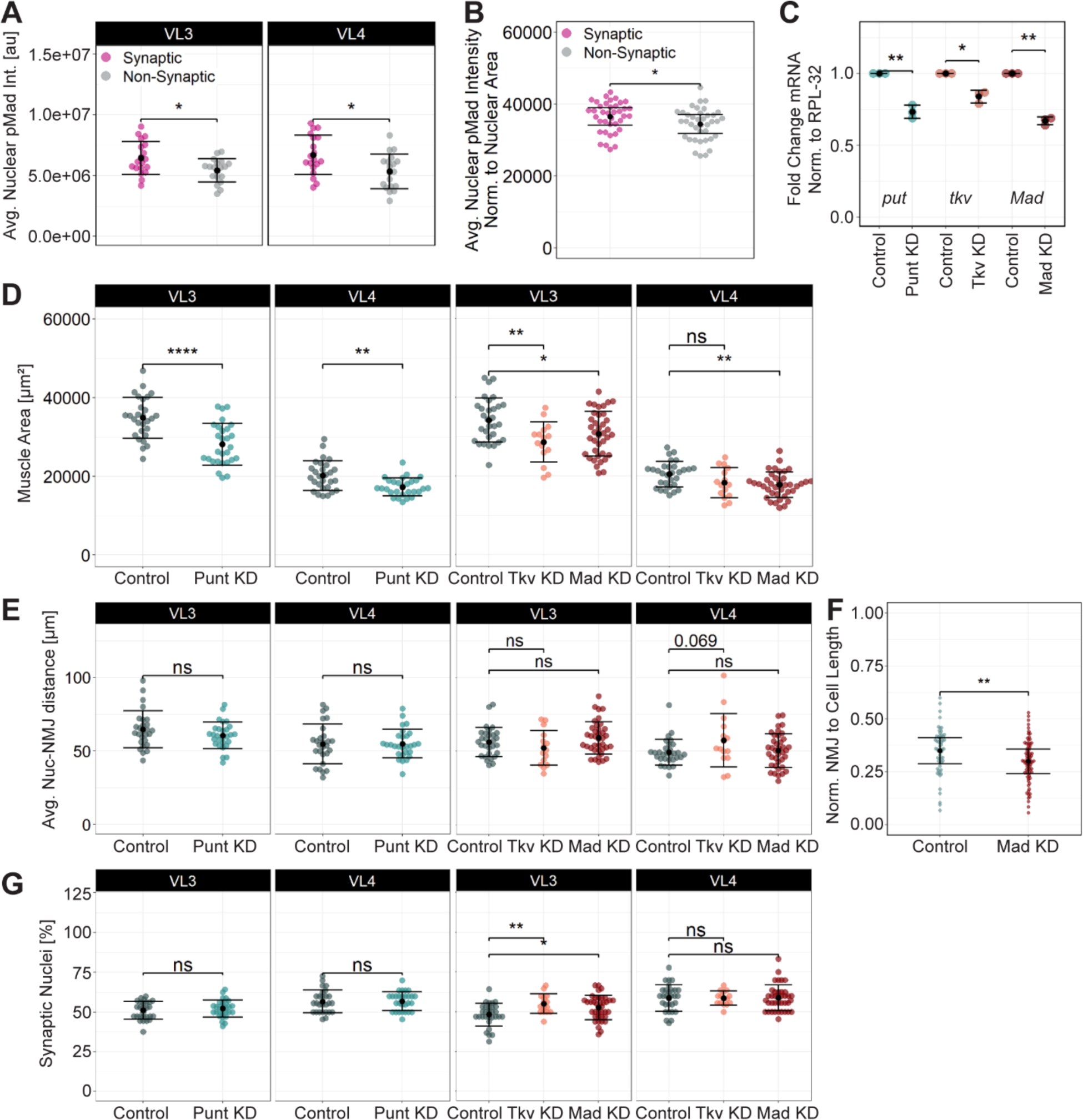
Muscle cell size and myonuclear proximity to the NMJ following KD of BMP signaling components. **(A)** Average nuclear pMad intensity in synaptic and non-synaptic myonuclei of VL3 and VL4 control muscles (*n =* 11 larvae, 40 VL3/4 muscle pairs). **(B)** Average nuclear pMad intensity normalized to nuclear area in synaptic and non-synaptic control myonuclei. **(C)** qPCR data showing Punt, Tkv and Mad KD (KD) efficiencies. Fold change in expression of *put*, *tkv*, and *Mad* mRNA normalized to housekeeping gene *rpl-32* (3 replicates; *put* and *tkv*: *n =* 10-15 larvae per genotype; *Mad*: 6-10 larvae per genotype). **(D)** VL3 and VL4 muscle area based on phalloidin labeling for control and Punt KD muscles (Control: *n =* 17 larvae, 28 VL3/VL4 muscle pairs; Punt KD: *n =* 17 larvae; 28 VL3/VL4 muscle pairs), and control, Thickveins, and Mad KD muscles (Control: *n =* 19 larvae, 30 VL3/VL4 muscle pairs; Tkv KD: *n =* 12 larvae, 15 VL3/VL4 muscle pairs; Mad KD: *n =* 20 larvae, 38 VL3/VL4 muscle pairs). **(E)** Average nuclear distance to the NMJ in VL muscles from Punt, Tkv, and Mad KD muscles and corresponding controls. **(F)** NMJ length normalized to muscle cell length in control and Mad KD muscles (Control: *n =* 19 larvae, 60 muscles; Mad KD: *n =* 20 larvae, 76 muscles). **(G)** Average percent of synaptic myonuclei in VL muscles from Punt, Tkv, and Mad KD muscles and corresponding controls. Data shown as mean ± SD for all plots, except in B and F where mean ± SEM is plotted (* p < 0.05, ** p < 0.01, **** p < 0.0001).

**Figure S3.**
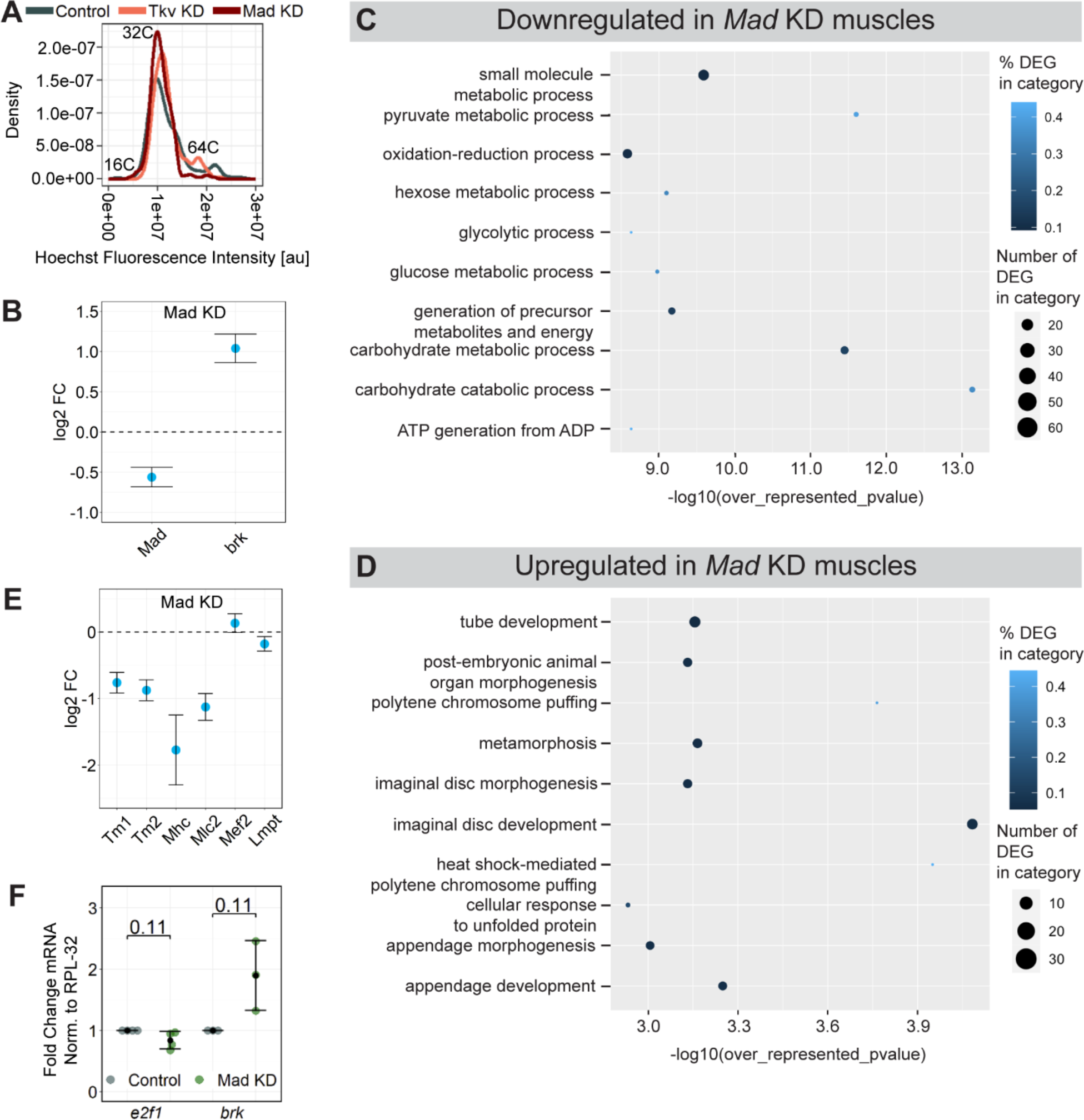
Genes and pathways associated with muscle growth are affected by KD of postsynaptic BMP signaling components. **(A)** A smooth density estimate plot of myonuclear Hoechst intensities in control, Tkv KD, and Mad KD muscles. Peaks indicate 16C, 32C, and 64C nuclei. **(B)** Fold change of *Mad* and *brk* mRNA in Mad KD compared to control muscle preps. **(C, D)** Top 10 **(C)** downregulated and **(D)** upregulated Gene Ontology (GO) gene sets for biological processes in Mad KD compared to control muscles. **(E)** Fold change of structural genes in Mad KD muscles. **(F)** qPCR data showing downregulation of *E2f1* and upregulation of *brk* in Mad KD muscles (*N =* 3-4 replicates; *n =* 6-12 larvae per genotype). RNA sequencing data was used for plots B, C, D, and E (*N =* 3 replicates, *n =* 6-10 larvae per genotype). Data shown as mean ± SD.

**Figure S4.**
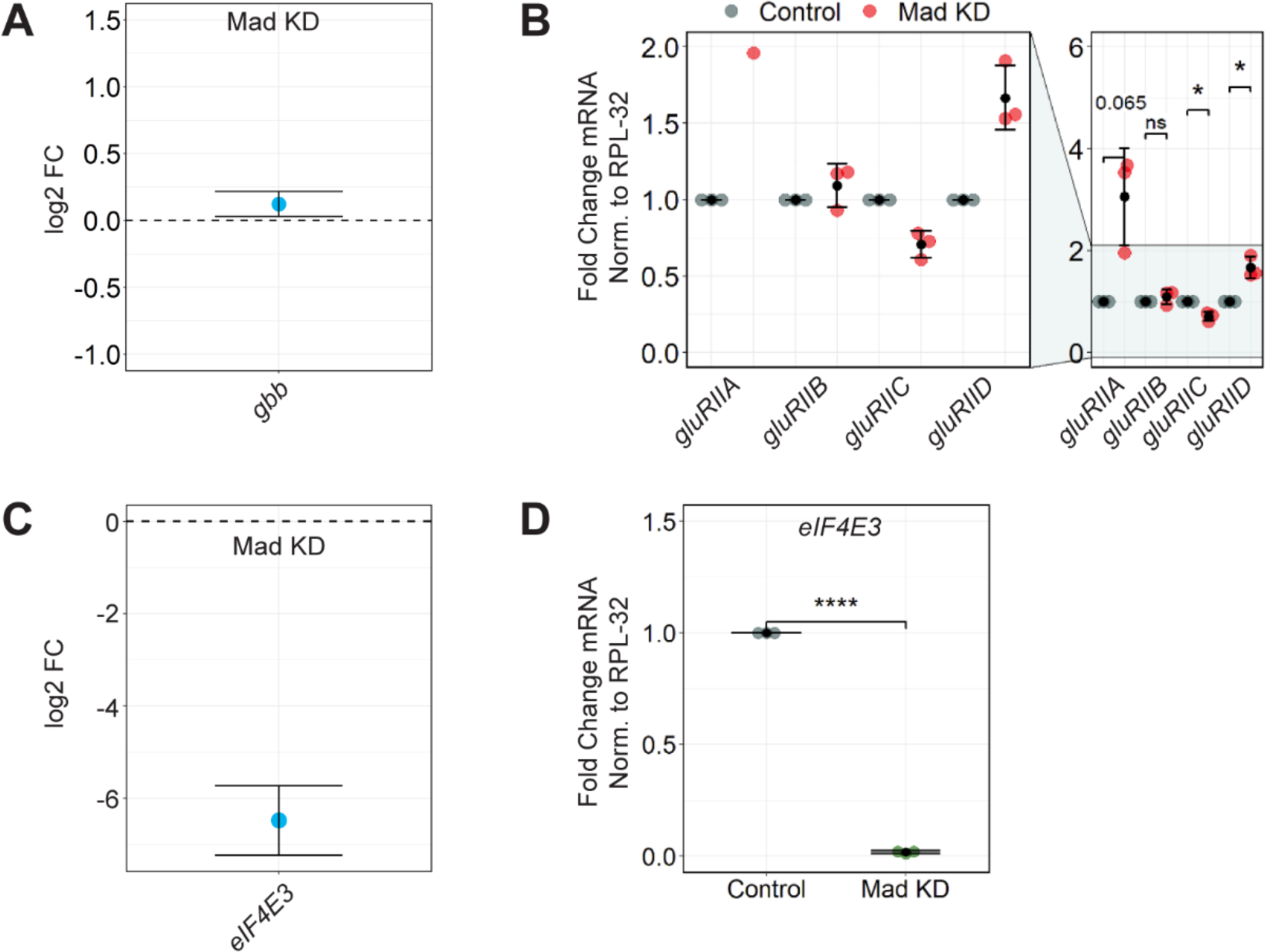
Expression of GluRII subunits and postsynaptic mRNA translation regulator eIF4E3, but not BMP ligand Gbb are dysregulated in Mad KD muscles. **(A)** Fold change of *gbb* in Mad KD compared to control muscle preps. **(B)** qPCR was conducted on GluRII subunits in control and Mad KD larval muscle preps to validate RNA sequencing results (*N =* 3 replicates; *n =* 10-14 larvae per genotype). **(C)** Fold change of *eIF4E3* in Mad KD compared to control muscles. **(D)** qPCR results of *eIF4E3* expression in control and Mad KD larval muscle preps (*N =* 3 replicates; *n =* 6-10 larvae per genotype). RNA sequencing data was used for plots A and C (*N =* 3 replicates, *n =* 6-10 larvae per genotype). Data shown as mean ± SD (* p < 0.05, **** p < 0.0001).

**Table S1.**
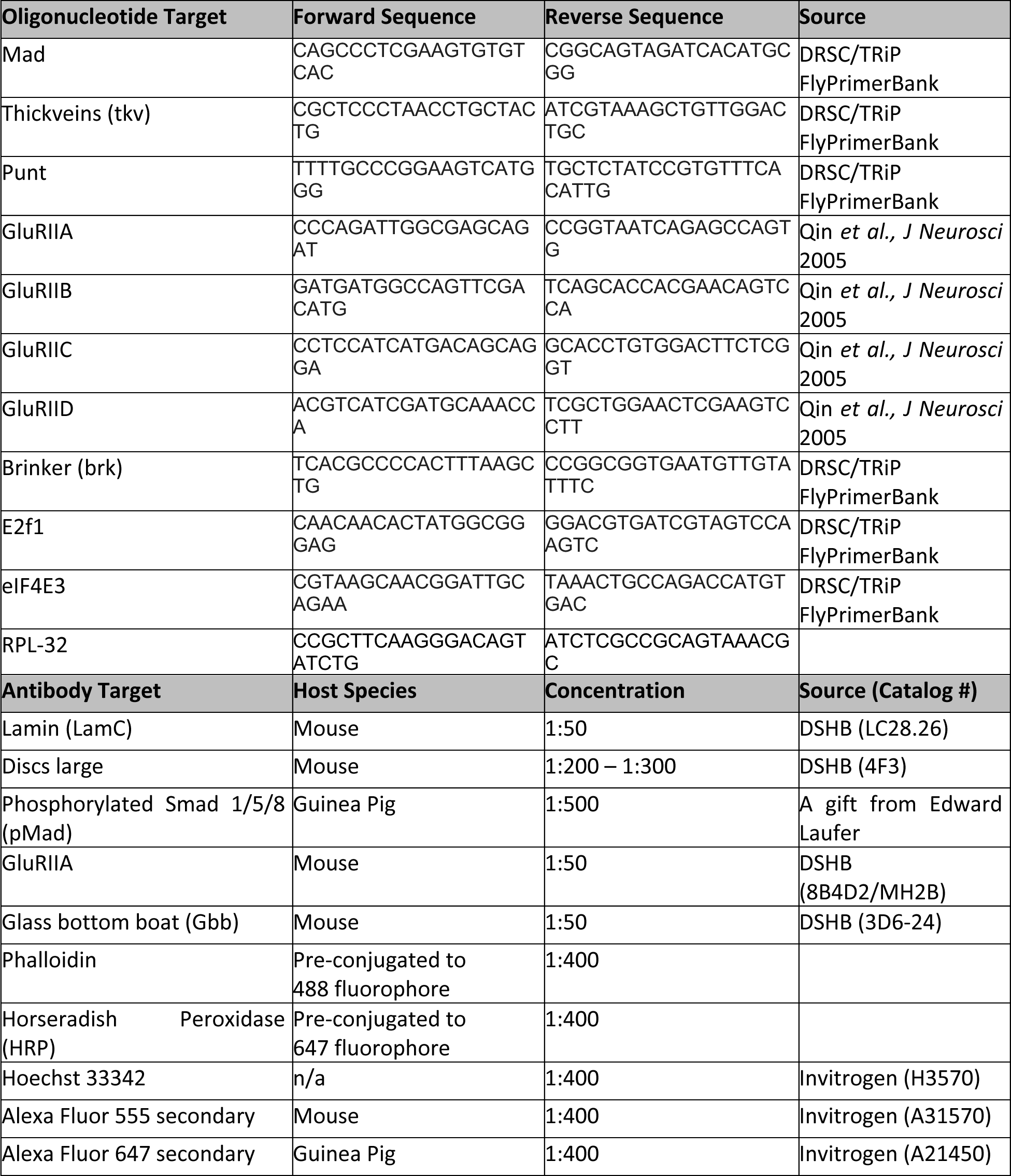
List of antibodies and oligonucleotides used in the study.

